# Behavioral, experiential, and physiological signatures of mind blanking

**DOI:** 10.1101/2024.02.11.579845

**Authors:** Esteban Munoz-Musat, Arthur Le Coz, Andrew W. Corcoran, Laouen Belloli, Lionel Naccache, Thomas Andrillon

## Abstract

Does being awake necessarily mean being conscious of something? This study investigates the phenomenon of Mind Blanking (MB), characterized by an "emptiness of mind", comparing it with Mind Wandering (MW) and On-task (ON) states. Using a sustained attention task and electroencephalogram monitoring, behavioral and neurophysiological signatures of MB were examined in 62 participants. MB exhibited a specific pattern of behavioral lapses, as well as decreased fast oscillatory activity and complexity over posterior electrodes compared to MW. Functional connectivity analyses revealed decreased long-range inter-areal connectivity during MB, compared to both ON and MW states. Event-related potentials with source reconstruction and temporal decoding techniques indicated a significant disruption in visual processing during MB, starting from 200 ms post stimulus and echoing into the late-stage of visual processing, suggesting a disruption of conscious access to sensory information during MB. EEG-derived markers allowed the prediction of mental states at the trial level, offering a finer view of conscious dynamics than subjective reports alone. Overall, these findings challenge the notion of the wakeful conscious mind as inherently content-oriented, suggesting that MB reflects genuine gaps in the stream of conscious thoughts, arising from disruptions in the generation or accessibility of thought content.

**SIGNIFICANCE STATEMENT:** Employing cutting-edge neurophysiological techniques on high-density electroencephalographic recordings, our study unveils unique neurophysiological markers of mind blanking, a phenomenon characterized by lapses in conscious content amidst the flow of wakeful consciousness. Distinguished from content-oriented states such as on-task and mind-wandering, mind blanking appears to be a distinct mental state. Furthermore, we demonstrate the feasibility of decoding consciousness dynamics solely from EEG features, transcending the limitations of intermittent subjective reports. Our findings thus not only provide a novel framework for investigating the stream of consciousness but also challenge the conventional notion that wakefulness invariably signifies being conscious of something.

## INTRODUCTION

> *“Consciousness, then, does not appear to itself chopped up in bits. Such words as ’chain’ or ’train’ do not describe it fitly as it presents itself in the first instance. It is nothing jointed; it flows. A ’river’ or a ’stream’ are the metaphors by which it is most naturally described. In talking of it hereafter let us call it the stream of thought, of consciousness, or of subjective life”* (William James, 1890).

In the former quote, William James pointed to two very intuitive aspects of our conscious experience^1^. First, consciousness seems *continuous* during wakefulness, without any “pauses” or breaks in the flow of contents of experience. Second, our conscious experience is dynamic and the origin of conscious content changes very frequently. Indeed, the stream of experiences can quickly shift between different external sources of information, but also turn inward towards internally generated task-unrelated thoughts, a phenomenon usually referred to as mind wandering (MW)^2,3^. Extensive research has shown that, in the brain, the transitions between task-related focus and MW involve modulations of activity of the Default-mode network (DMN)^4^ as well as changes in cortical dynamics with modulations of alpha activity^5,6^ and increases in slower rhythms^7^. However, a rigid separation between MW and task-focused states has been challenged by evidence emphasizing the role of context in shaping the neural correlates of MW^8,9^. Recent approaches, sometimes leveraging more naturalistic paradigms, focus instead on the experiential features of subjective experience: e.g., what kind of information is being processed and how (e.g., level of detail, engagement)^10,11^. For example, MW research has investigated the distinctions between different MW subtypes, stressing the importance of considering the meta-awareness (i.e., were participants aware of their MW) or voluntariness (i.e., did they mind-wander on purpose?) dimensions of MW^12^. This perspective frames mind-wandering and task-focused states as points along a continuum within a multidimensional phenomenological space^13,14^.

More recently, a new “mind state” has been described, challenging the idea of a continuous stream of conscious contents during wakefulness: the state of mind blanking (MB). MB is described as a waking state that is either spontaneous or intentional, during which a subject does not report any mental content, but rather the feeling of an empty mind^15^. Previous studies have shown that MB is reported about 14-18% of times during focused tasks^15,16^ and about 6% of times during resting^17^. Furthermore, MB has been associated with a specific behavioral outcomes, distinct from both MW and task-focused states^7^. The exact nature of mind blanking is still a matter of debate^18,19^ and only a few studies have attempted to investigate its neural correlates^20^. Some have reported a widespread deactivation of thalamic and cortical brain regions during spontaneous MB^21^ and a more focal deactivation of the superior and inferior frontal gyri and hippocampus during intentional MB, whereas anterior cingulate cortex activation seemed to increase^22^. Others have reported an increase of functional connectivity, as measured by phase coherence metrics, during MB^23^. Finally, MB has been linked to the occurrence of sleep-like slow waves in scalp EEG^7^.

Two pressing questions regarding MB remain. First, it remains unclear whether MW and MB correspond indeed to different subjective and neurophysiological states^24,25^ or if they can be traced back to common underlying physiological causes expressed in varying degrees^26^. Recent evidence suggests that both MW and MB could be explained by local sleep phenomena in the brain, but with different regional distributions^7^. Other accounts suggest that MB could arise from a specific pattern of neural (de)activations or functional connectivity, distinguishable from both task-focused and MW states^21,23^. To better understand the true nature of MB in respect to both task-focused and MW states, the neural distinctions of these mind states, if any, need to be further investigated^27,28^.

Second, the place of MB in the hierarchy of consciousness states needs to be clarified. Traditionally, a clear distinction was made between states of unconsciousness (e.g., coma, deep N3 sleep) and states of consciousness (e.g., normal Wakefulness, REM sleep). States of consciousness (i.e., being conscious) classically imply the existence of conscious contents (i.e., being conscious of something) whereas states of unconsciousness imply the absence of such contents. Recent work suggests that the frontier between conscious and unconscious states is not as clear-cut. Indeed, conscious experiences (dreams) are often reported during consolidated non-REM sleep^29,30^; and transient behavioral as well as electrophysiological signs of conscious processing have been recently reported during sleep including N3 sleep^31–33^. These results suggest an intriguing possibility: could the reverse phenomenon - transient absence of conscious content during wakefulness- exist^27^? If so, could MB be a potential candidate for such a particular state?

Some theoretical accounts of consciousness align with the possibility of gaps in the stream of conscious contents. For example, the Global Neuronal Workspace Theory of Consciousness (GNWT)^34,35^ suggests that conscious access (and hence, reportability) to a given representation depends on the late, sustained and global ‘ignition’ of fronto-parietal associative areas. According to this view, consciousness is a discrete phenomenon, and windows of unconsciousness during Wakefulness could result from a gap between two ignitions. The Integration Information Theory (IIT) posits that consciousness would dissolve whenever the brain’s capacity to integrate information breaks down^36^. Changes in cortical dynamics towards less integration may correspond to gaps in the stream of consciousness, much like the mirror phenomenon, local activations within sleep, is linked to dreaming^30^. The parallel with sleep and dreaming research is particularly inspiring, since extensive research has shown that (macro) unconscious states (N3 sleep, deep anesthesia, coma, the vegetative state) are characterized by : i) a decrease in neural complexity and fast neural oscillations^37^–40; ii) a breakdown of information sharing between distant cortical areas^41–44^, particularly in the delta and theta frequency bands^45,46^ ; iii) impaired neural responses and representations to external stimuli, particularly during the late-stage of information processing (>300ms post-stimulus)^39,47–51^. These previous studies provide candidate EEG markers to explore the signature of MB.

Here, we present compelling evidence that MB is a separate mental state, distinct from MW and task-focused states at the phenomenological, behavioral and neurophysiological levels. To provide an exhaustive characterization of the neurophysiological fingerprint of MB, we relied on previous work exploring consciousness states and contents^52^. In particular we capitalized on the use of markers tracking both background neural dynamics (complexity, information sharing) and the processing of external information (ERPs, temporal decoding) in the sensor and source space. Based on these signatures, we argue that MB could represent a failure of conscious access mechanisms, resulting in a genuine gap in the stream of conscious contents.

## RESULTS

### Different behavioral signatures between Mind Blanking and Mind Wandering

In these two studies, 62 healthy participants performed a modified sustained attention to response task (SART), with digits and faces as stimuli presented in separate blocks, while high-density electro-encephalogram (hdEEG) was recorded continuously. After each stimulus presentation, participants had to press a button (Go trials), or refrain from doing so whenever a “3” or a smiling face was presented (No Go trials; 1 out of 9). Stimulus onset asynchrony (SOA) varied randomly (from 750-1250ms) with a null inter-stimulus interval, and participants’ mental state was probed at random intervals (every 40-70s, uniform random jitter). Participants were instructed to report their attentional focus ‘just before the interruption’ by selecting one of four options: (1) ‘task-focused’ (ON), (2) ‘off-task’ (mind-wandering, MW), (3) ‘mind-blanking’ (MB), or (4) ‘don’t remember’. Since the fourth option accounted for only 2.6% of all probes (i.e., less than two probes per participant on average) and given that previous studies did not consistently distinguish between these options^53^, we merged third and fourth options as MB in all analyses (Figure 1A-B).

**Figure 1:**
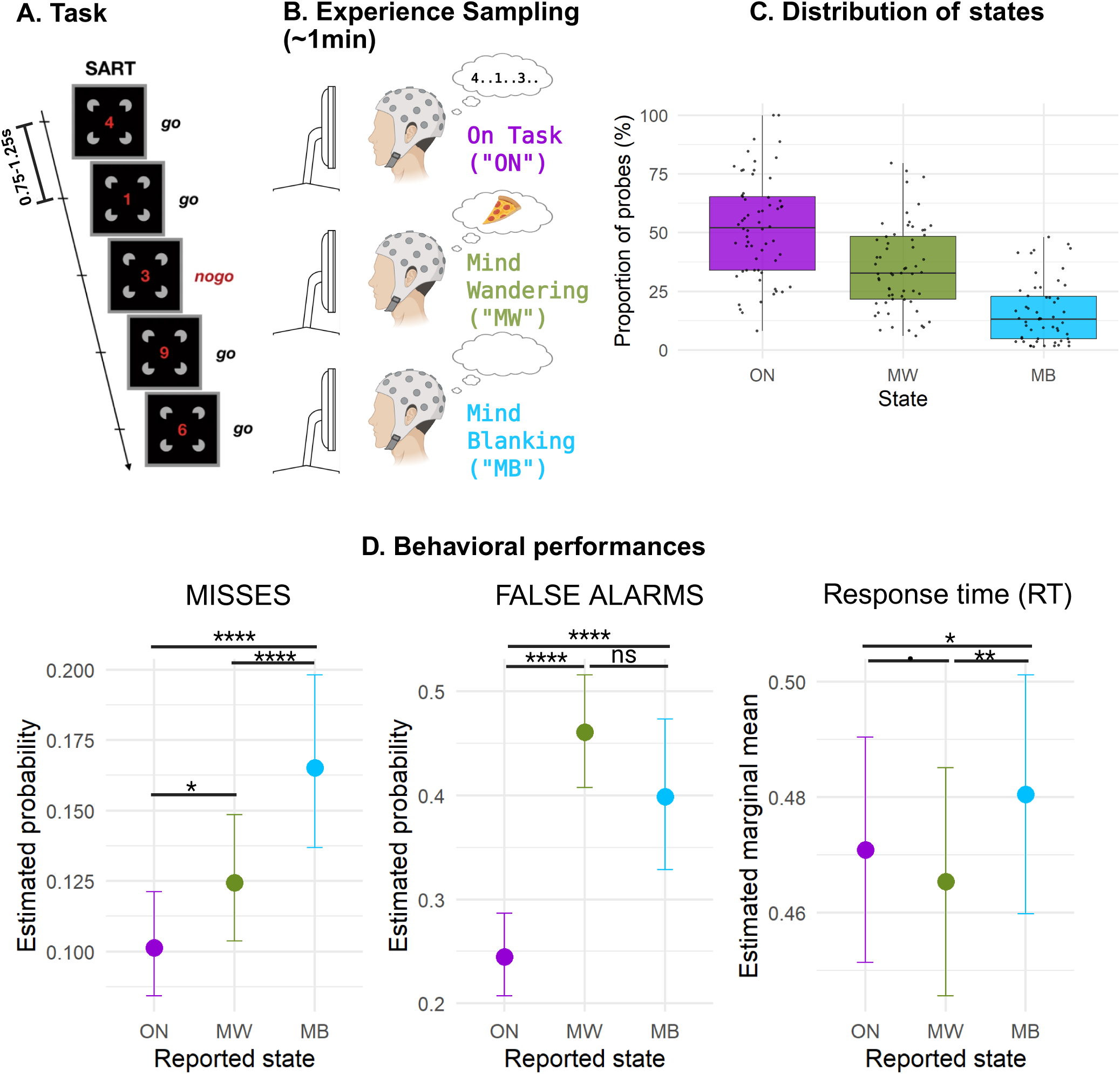
Experimental paradigm and behavioral results. Panels A and B: Experimental paradigm. 62 healthy participants performed a sustained attention to response task (SART), with digits and faces as stimuli, while high-density electro-encephalogram (EEG) was recorded continuously. After each stimulus presentation, participants had to press a button (Go trials), or refrain from doing so, whenever a “3” or a smiling face was presented (No Go trials; 1 out of 9). SOA varied randomly (750-1250ms) with a null ISI (panel A). Participants’ mental state was probed at random intervals (every 40-70s, uniform random jitter). They had to report their attentional focus by selecting one of four options: (1) ‘task-focused’ (ON), (2) ‘off-task’ (mind-wandering, MW), (3) ‘mind-blanking’ (MB) (panel B). Panel C: proportion of probes in function of the reported mind-state. Proportions were computed at subject level (individual dots) and the distributions at group level are represented by a boxplot (the boundaries of the boxes represent first and third quartiles (Q1 and Q3 respectively), mid-line represents the medium and the whiskers depict Q1-1.5*IQR and Q3*1.5*IQR). Panel D: Behavioral performances in function of the reported mind-state. Misses, False Alarms and Response Times (RT) were computed for the trials between the 5s before probe-onsets. (Binomial) linear mixed models were computed with *mind-state* as the main explanatory factor, and subject ID/dataset as a random intercepts. We found fine-grained modulations of performance in function of mind-state, with distinct behavioral profiles for MW (faster RTs) and MB (slower RTs, more misses). The statistical bars and stars represent the pairwise comparisons between mind-states (FDR corrected). The detailed statistical results can be found on Table 1. ****: FDR corrected p-value<0.0001 **: FDR corrected p-value<0.01 *: FDR corrected p-value<0.05 . : FDR corrected p-value<0.1 n.s. FDR corrected p-value>0.1

Participants performed this task with a high accuracy (mean accuracy of 86.5 ± 0.5% (SEM)) and with a relatively low rate of Misses (i.e., errors in Go trials, 10.4 ± 0.5%). Errors in No Go trials (False alarms [FA]) were frequent (36.7 ± 1.5%). At the subjective level, participants reported being focused on the task (ON) in only 52 ± 2.7% of the probes and otherwise reported MW (35 ± 2.3%) or MB (16 ± 1.8%) (Figure 1C). The distribution of the ON, MW and MB mind-states was not uniform across participants (see Figure 1C), with large inter-individual differences: in particular 6 participants (9.6%) did not report any MB experience but reported MW experience, 2 participants (3.2%) reported no MB and no MW and no participants reported MB without MW.

In order to detect fine-grained modulations of behavioral performance according to the participant’s mind-state, we labeled all trials presented up to 5 seconds before each probe as corresponding to the probed state (e.g., after an ‘ON’ response, all previous trials that occurred during the previous 5 seconds were labeled as ‘ON’ trials). As previously reported in a smaller subgroup of participants (40% of the present cohort)^7^, we found a significant main effect of *mind-state* on FA, misses and RTs (see Table S1). Crucially, pairwise comparisons between mind-states revealed different behavioral profiles for MW and MB: while both were characterized by more FA and misses than during ON, FA were equally frequent during MW and MB (with a non-significant trend for more FA during MW trials) and misses were more frequent during MB (i.e. MB>MW>ON) (see Table S1 and Figure 1D). Response times (RTs) were *slower* for MB (compared to both ON and MW), and *faster* for MW (compared to MB and ON; significant difference for MW vs MB and statistical trend (FDR corrected p-value 0.1) for MW vs ON) (see Table S1 and figure 1D). In sum, while MW seemed characterized by an “impulsive profile” (more FA and faster RTs), MB presented with an “absent profile” (more misses and slower RTs), which confirms our previous findings obtained in one of the two datasets.

### ‘Front-Back’ dissociation between Mind Blanking and Mind Wandering

We next investigated neurophysiological correlates of the mind-states (ON, MW and MB) from EEG recorded during the 5s preceding each probe onset. We computed three classes of neural markers: 1) spectral markers (normalized power spectral densities (PSD) in the delta, theta, alpha, beta and gamma bands); 2) complexity markers^39,54^ (Kolmogorov Complexity (KC) and Sample Entropy (SE)) and 3) connectivity markers *(weighted symbolic mutual information*^45^ (wSMI) in the delta, theta and alpha bands). These markers (and other similar measures) have been previously shown to track modifications of cognitive and consciousness state in a large range of conditions, including disorders of consciousness (DoC)^39,45,55^, sleep^46,56,57^, hypnosis^58^ and meditation^59^.

All studied neural markers were modulated by the participants’ mind-state. A significant main effect of *mind-state* and of *electrode location* was found for the wSMI in the delta and theta frequency bands, and a significant effect of these two main factors (mind-state, electrode location) and their interaction was found for all other markers (wSMI alpha; normalised PSD delta, theta, alpha, beta, gamma; sample entropy and Kolmogorov complexity) using linear mixed models with mind-state, electrode location and their interaction as main factors. Subject ID and dataset (see Methods) were introduced as random intercepts (see Table S2).

Spectral and complexity measures revealed different profiles for MW and MB: compared to ON, MW was associated with both an increase in fast oscillatory activity and complexity over central electrodes, and with a decrease of these metrics over fronto-polar and posterior electrodes. As for MB, compared to ON, it was associated with an increase in fast oscillatory activity and complexity over frontal and fronto-central electrodes, and with a decrease in these metrics over centro-posterior and posterior electrodes. The direct contrast between MB and MW revealed a front vs. back dissociation: while frontal and fronto-central electrodes showed faster oscillatory activity and higher complexity in MB than in MW, centro-posterior and posterior electrodes showed an opposite difference (see Figure 2A,C).

**Figure 2:**
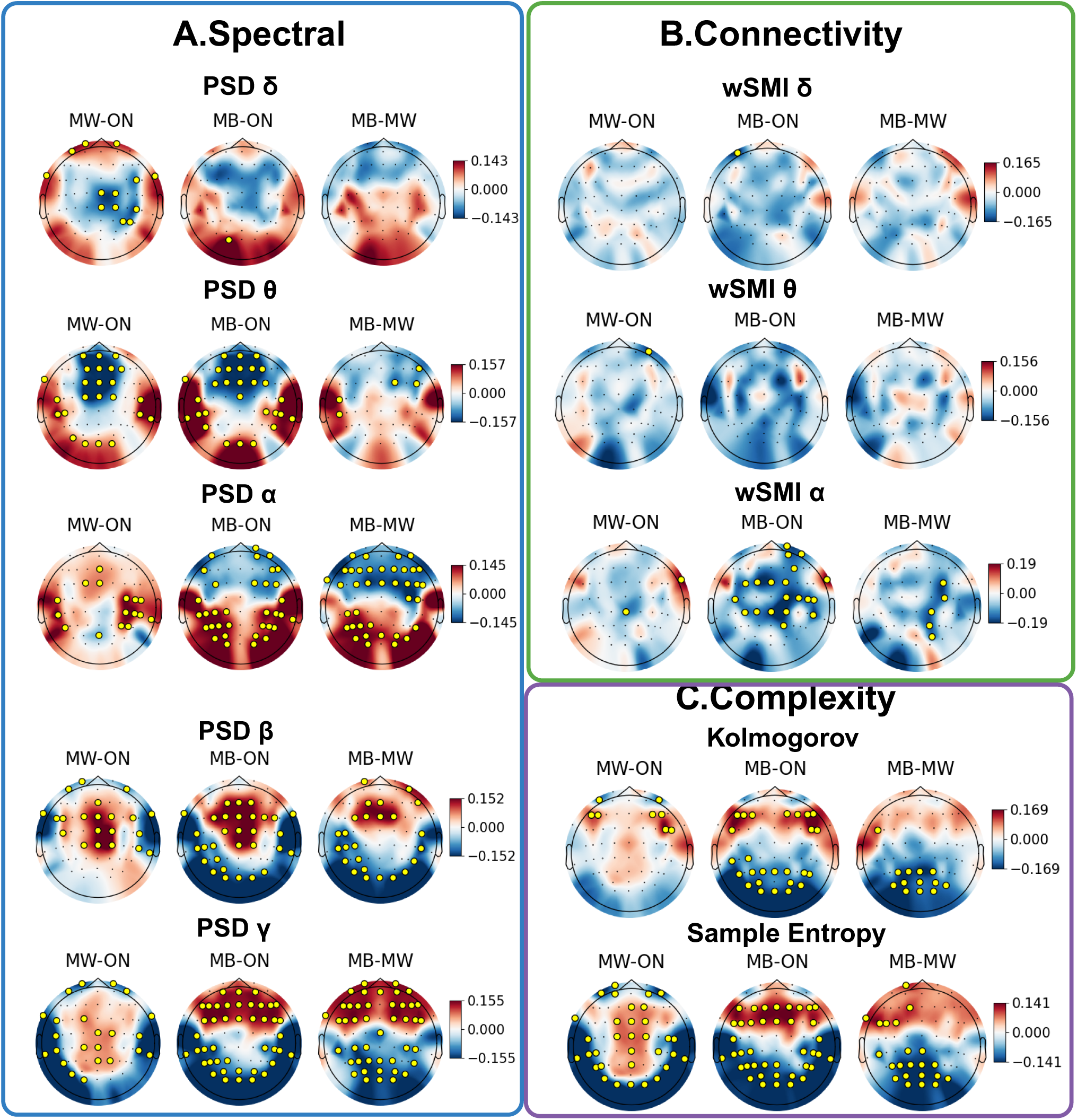
EEG-based neurophysiological markers differentiate MB from ON and MW. Spectral (A), Connectivity (B) and Complexity (C) results. Each subplot represents the pairwise statistical contrast between two states (left: MW - ON; middle: MB - ON; right: MB - MW), at sensor level (topographical representation of the scalp). Model estimates for the contrast between states were computed for each electrode, and locations with statistically significant differences (FDR corrected p-value<0.05) are depicted with a golden circle. We observed a front-back dissociation between MW and MB for spectral and complexity measures, and a progressive breakdown of functional connectivity going from ON to MB (ON>MW>MB). The ANOVA tables summarizing the main factor’s effects (linear mixed models with mind-state, electrode/connection and their interaction as main factors, and subject ID/dataset as a random intercepts) for each EEG metric can be found in Table 2.

Mind-state can be modulated by arousal, with MW and MB frequency increasing in moments of hypovigilance^7,60^. Therefore, some of the observed changes in neural markers could be explained by a decrease in alertness. To ensure that our results couldn’t be *exclusively* explained by this arousal factor, we conducted a verification analysis, by including in our statistical model the vigilance score as a covariate (4-point scale, 1 = Extremely Sleepy, 4 = Extremely Alert), as well as the interaction between electrode location and vigilance score. The observed results for spectral and complexity measures were nearly identical to those previously presented, with the persistence of a clear front-back dissociation between MW and MB. See supplementary Figure S1A,C.

### Breakdown of long-range information sharing during Mind Blanking

Functional connectivity analyses revealed a significant modulation by *mind-state* of the wSMI metric in frequency bands of interest (delta, theta and alpha) (see Table S2). Post-hoc topographical analyses in sensor space revealed a progressive breakdown of connectivity from ON to MW, and then from MW to MB, particularly in the alpha band (see Figure 2B). As for spectral and complexity measures, the inclusion of vigilance score as a covariate in statistical models resulted in very similar results (see Figure S1B). To better characterize these connectivity modulations in function of mind state, we reconstructed the cortical sources of the EEG signal (N=68 cortical sources) at the trial level and computed the information metric wSMI (at different frequency bands) between each pair of sources (n=2278 connections). These sources were further grouped in 10 pairs (right and left) of 5 ROIs, according to the Desikan-Killiany atlas (frontal, limbic, temporal, parietal and occipital), obtaining thus 45 averaged connections between ROIs. Since previous studies reported modifications of coherence-based metrics during MB, we also computed in source space the Phase Locking Value (PLV), for comparison. The statistical analysis revealed a main effect of *mind-state* for all computed connectivity metrics (linear mixed model with mind-state, connection and their interaction as main factors; subject ID and dataset were introduced as random intercepts; see Table S3). As reported in a previous study^23^, the coherence-based metric PLV showed an increase in inter-areal connectivity during MB, compared to both MW and ON states (see Figure 3, top panel). By contrast, the information-based metric wSMI showed the reverse pattern: for all computed frequency bands (delta, theta and alpha), a significant reduction in inter-areal connectivity was observed during MB, compared to both ON and MW (except for the connectivity in the delta band between right frontal and right occipital areas, which increased) (see Figure 3). This disruption of inter-areal connectivity during MB concerned mainly parietal areas, and in particular fronto-parietal connectivity.

**Figure 3:**
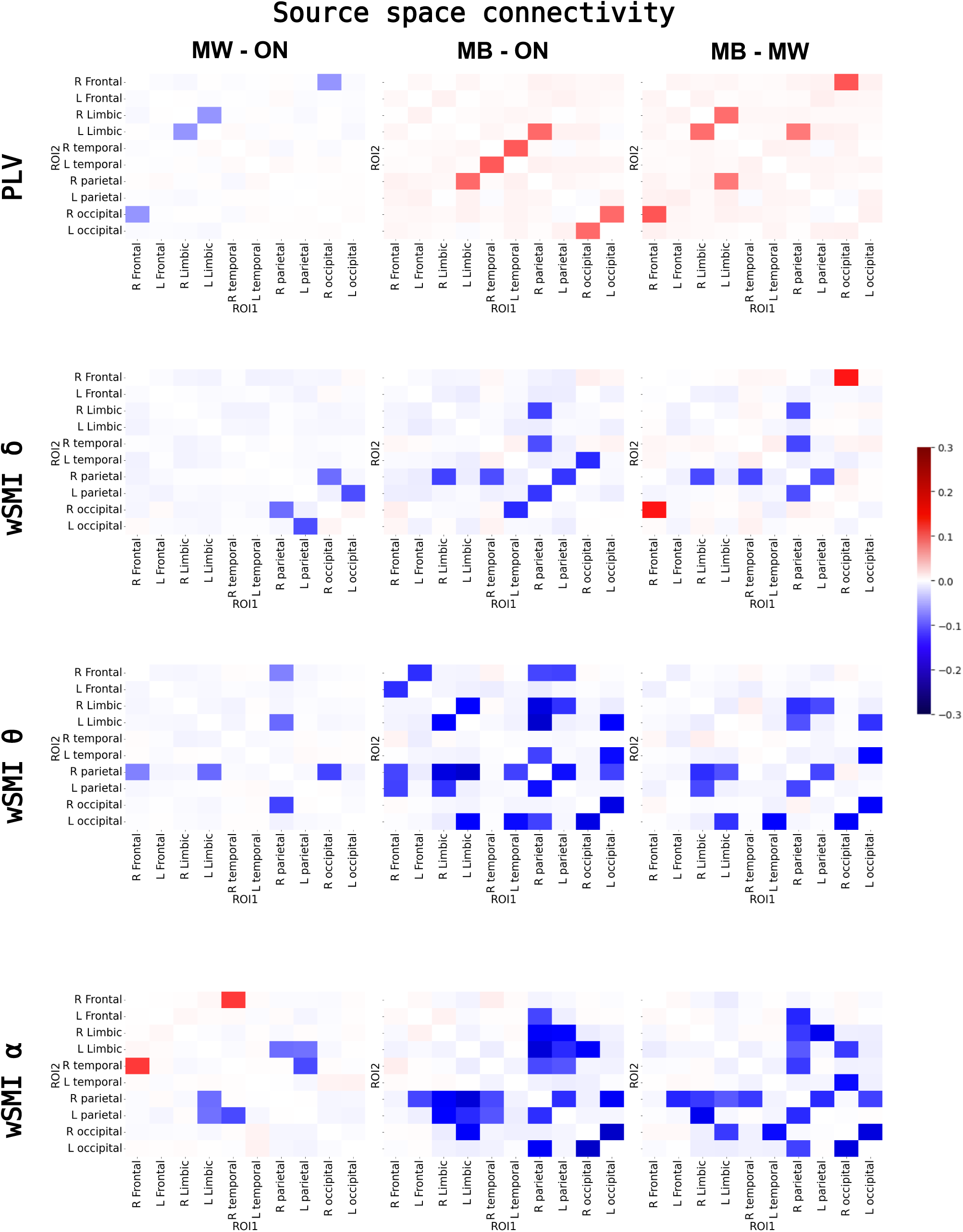
Increased phase synchrony with reduced information sharing during MB. The PLV (top) and the wSMI in different frequency bands were computed at the source level. Each square-matrix represents the contrast in connectivity, computed in source space, between two mind-states, for each pair of regions of interest (ROIs). Only significantly different connections (FDR corrected p-value<0.05) are highlighted, the other ones are masked. ANOVA tables for the statistical models can be found in Table 3. We observed a dissociation between the PLV and wSMI, with increased PLV (phase synchrony) and decreased wSMI (information sharing) between distant cortical areas (in particular fronta-parietal) during MB, as compared to ON and MW. L: delta band; θ: theta band; α: alpha band wSMI: weighted symbolic mutual information PLV: Phase Locking Value R: right; L: left

The inclusion of the vigilance score in the statistical models led to results largely consistent with those previously presented for the PLV and the wSMI in the theta and alpha bands; for the wSMI in the delta band, the results were more nuanced, revealing both increases and decreases of long-range connectivity during MB (see Figure S2).

As it will be discussed later, the contrast between the PLV and the wSMI results could reflect the relative implication of linear vs. non-linear interactions between cortical regions during these different mind states, non-linear interactions being best captured by the wSMI (see discussion).

As an interim conclusion, these neurophysiological findings confirmed and extended our behavioral results, by showing that MB is associated with a specific brain state distinct from MW. In addition, the significant disruption of long-range connectivity revealed by the wSMI metric during MB suggests a marked impairment of information sharing between cortical areas during this mind-state, as systematically observed during unconscious states.

### Disruption of the brain processing of external stimuli during Mind Blanking

The processing of external information is very dependent upon the background brain activity and associated mind state, and foremost the state of consciousness. Previous research has shown that early and intermediate cortical processing is preserved in classically unconscious states such as non-REM sleep^50,61–65^ or coma^66–68^ (having a prognostic value in this last case). By contrast, late cortical processing seems to be associated with conscious states^31,47,48,50^ and conscious processing of external information^69–72^. With this in mind, we decided to probe the differential neural fate of the presented visual stimuli according to the reported mind-state.

First, we conducted a single-trial multi-level analysis of event-related potentials (ERPs) in sensor space (see Methods for details and Table S4 for the results of the statistical model). Early processing of visual stimuli was extremely similar across ON, MW and MB mind-states. In particular, a similar posterior positivity peaking around 100-150 ms after stimulus presentation, and corresponding to the P1 component^73^, was present in all 3 conditions (see Figure 4A). Visually, this P1 component seemed more pronounced in ON and MW states than in MB, but very few electrodes showed a significant effect (≤3). No significant differences were observed between MW and ON during the early time-window (see Figure 4B). Crucially, stimulus processing diverged massively as a function of mind-state during the late time-window (>350ms). In ON trials, a classical P3b component (central positivity) spanned roughly from 400 to 650 ms after stimulus onset. In MW trials, this P3b pattern was still observable, but with a longer latency, a reduced maximal amplitude and a shorter duration. Finally, in this late time-window, no clear P3b response was observable in MB trials; amplitudes of MB ERPs were significantly decreased as compared to ON and MW, with only a reduced and left-lateralized response, possibly related to motor responses (Figure 4A-B). As a control, we also conducted the previously described analysis independently for each stimulus type (faces vs digits). ERP profiles show differences in the timing of early and late components yet, for digits and faces, MB trials were characterised by a disruption of late components compared to ON trials (see Figure S3).

**Figure 4:**
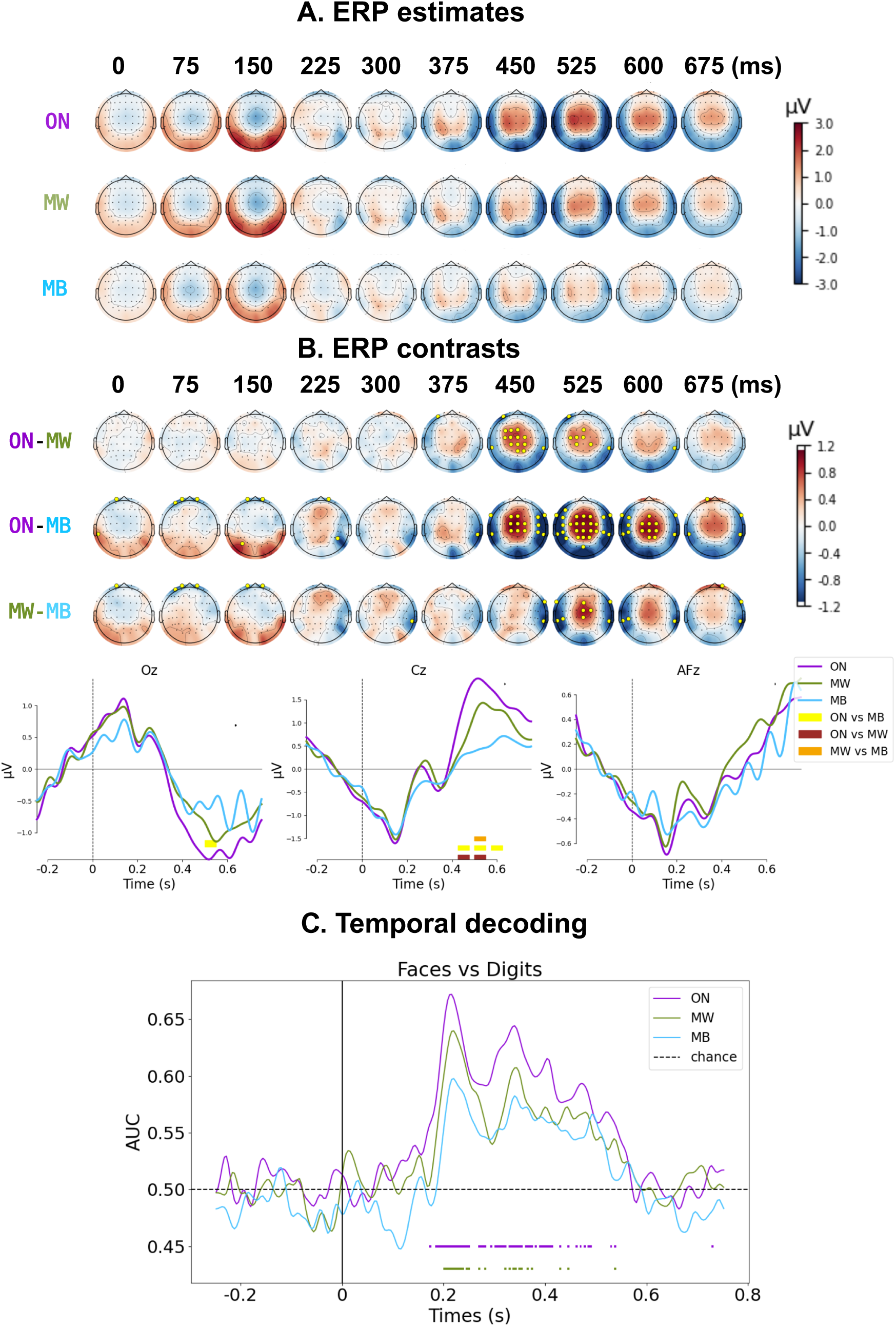
Stimulus-induced activity analyses. Panels A and B: Topographical scalp representation of the ERP estimates for each mind state (A) and ERP contrasts between mind-states (B), derived from the statistical model (single-trial multi-level analysis with mind-state, EEG channel and their interaction as main factors, and subject ID/dataset as random intercepts). Marked electrodes in panel B (golden circles) are the ones presenting a statistically significant difference between conditions (FDR corrected p-value<0.05). Panel B bottom presents the grand-average ERP time-series for each mind-state (ON: violet; MW: green; MB: blue) for 3 electrodes of interest (Oz, Cz and AFz). Time-points with statistically significant differences between conditions (FDR corrected p-value<0.05) are highlighted (ON vs MW: brown; ON vs MB: yellow; MW vs MB: orange). Given that trials follow one another in close succession, it is normal for the baseline activity to be influenced by the activity of the previous trials. Panel C: Temporal decoding of stimulus category. A linear classifier was trained and then tested (via cross validation) at each time point to distinguish stimulus category (Faces vs Digits), allowing the tracking of the temporal dynamics of neural representations, independently for each mind-state. Note that the number of trials were identical between mind-states for this analysis. Classifiers’ performances were compared to chance-level performance (see Methods), independently for each mind-state, and statistically significant time-points were represented by colored horizontal bars. While both early (<300ms) and late (>300ms) time-points with significant decoding were observed for ON and MW, no significant decoding was observed for MB trials.

While univariate methods such as ERPs provide with important information about the amplitude and timing of cortical activations in response to external stimuli, multivariate decoding methods allow for a more subtle analysis, and provide information at the level of neural representations. We wanted to answer the following question: does brain activity encode relevant information about the presented stimuli, and how these neural representations vary as a function of the reported mind-state? To do so, we tested whether we could decode stimulus type (Faces vs. Digits) from brain responses using multivariate pattern analysis (MPVA), with the Temporal Decoding method^49,74^. Briefly, this type of analysis consists in using a subset of the data to train a linear classifier at each time-point to differentiate trials where a *Face* was presented from trials where a *Digit* was presented, and then testing its performance, independently at each time-point, in a different subset of the data. We ensured a comparable number of trials between mind-states by randomly subsampling trials from each mind-state. Then, we applied the Temporal Decoding method independently for each mind state, and statistically compared the obtained performance scores (ROC AUC) with a dummy random distribution centered around 0.5 (chance level performance). Results can be found in Figure 4C. For ON trials, we found several time-windows with significant decoding compared to chance level performance. The first one, around 200ms, coincided with a sharp decoding peak (AUC>0.65). The other ones, spanning from 250ms to 580ms, corresponded to a sustained period of decoding. For MW trials, we found a similar initial peak around 200ms; however, later time-windows with significant decoding were sparser, with a period around 350ms, and then a few significant time-points after 400ms. Crucially, we didn’t observe any statistically significant decoding for MB trials.

Given these results, we reconstructed the cortical sources of the EEG signals at trial level, and compared the post-stimulus activations (time-window: 0-700ms) to the activations during the baseline period (-250 - 0 ms). For both ON and MW trials, we observed a significant activation of bilateral occipital visual regions (around 200-300 ms post-stimulus), followed by an activation of regions along the ventral and dorsal visual streams (300-600ms), reaching some frontal areas (around 500ms post stimulus). Responses in MW trials were smaller, delayed and less spatially extended compared to ON. Importantly, for MB trials, we observed only sparse activations, mostly left-sided (contralateral to motor responses), with no clear involvement of the dorsal or ventral visual streams (see Figure 5).

**Figure 5:**
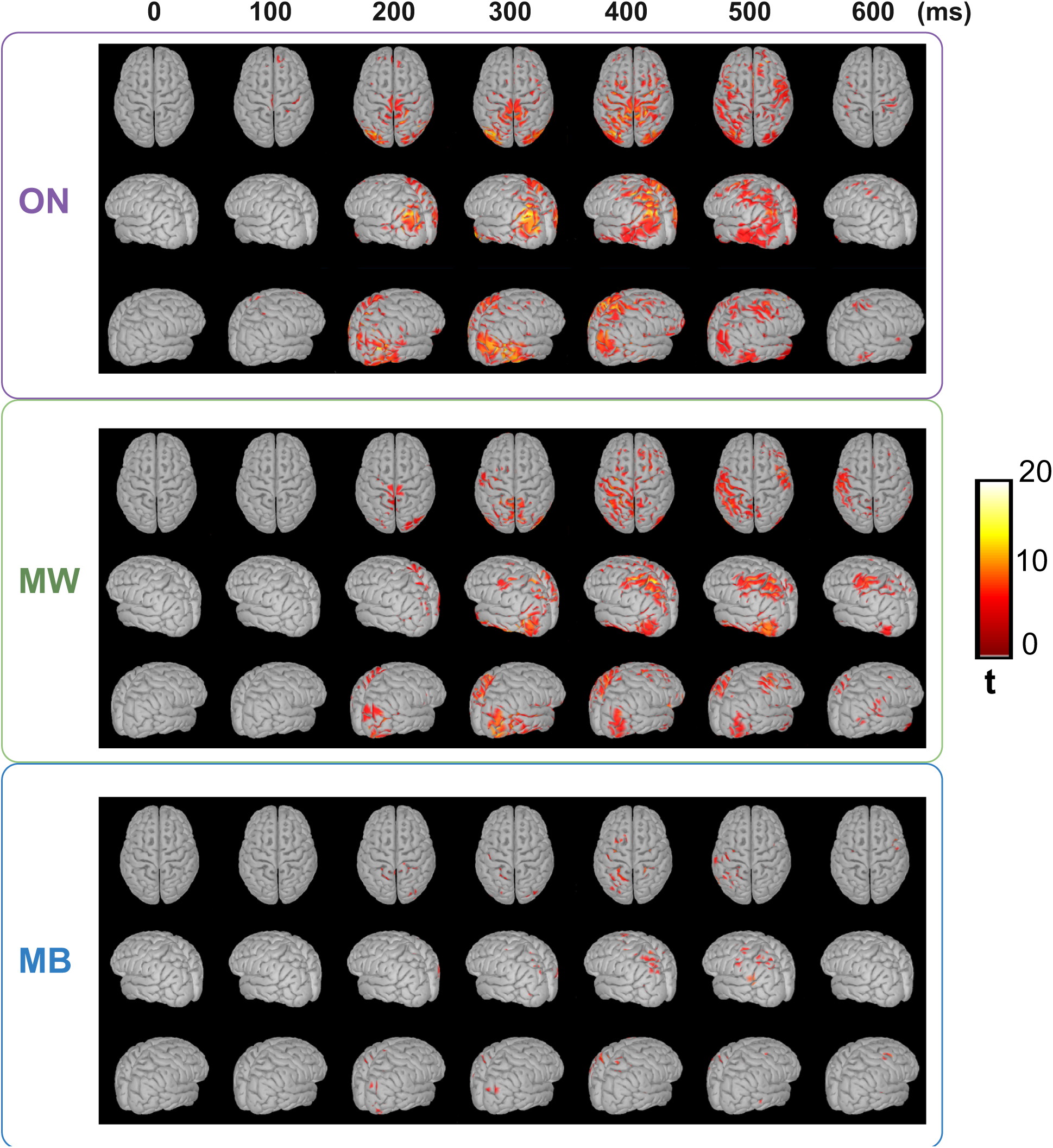
Source reconstruction of stimulus-induced activity. Source reconstruction at trial level was performed independently for each mind state, and a t-test against the baseline (-0.25 to 0s relative to stimulus onset) was performed. Only statistically significant modulations of activity compared to the baseline (FDR corrected p-value<0.05, corrected for multiple comparisons across time, space and frequencies) are highlighted. While significant activations of visual streams (dorsal and ventral) were observable for both ON and MW conditions, starting from 200ms post-stimulus, this was not observed during MB trials.

In sum, while the essential processing steps of visual information seem preserved during MW compared to ON-task, these processes appear significantly disrupted during MB, particularly the late responses usually associated with conscious access.

### Trial-by-trial prediction of mind-state based on EEG neural features

The exploration of the dynamics of consciousness is limited by the reliance on the discrete sampling of experience through mind probes and subjective reports. To circumvent this snapshot approach, we set out to predict participants’ mind-states on a trial-by-trial basis using neural features. We gathered all the previously presented EEG markers from stimulus-centered epochs (-0.25 to 0.75 s relative to stimulus onset), for all trials. We then trained and cross-validated, independently for each subject, machine learning classifiers (Random Forest), using exclusively trials within the 5 seconds preceding probe onsets (since we can label these trials as “ground truth” based on subjective reports). Importantly, during the cross-validation procedure, we grouped trials by experimental block (6 blocks for each participant) to avoid overfitting due to temporal proximity between trials (see Methods for more details). In multiclass classification (ON vs MW vs MB), the median classifier’s balanced accuracy score across participants was of 46%, significantly above chance level performance (median chance level (500 permutations): 37%, FDR corrected p-value=9.10^-6^, Wilcoxon rank-sum test, see Methods for more details ) (Figure 6A; Table S5 and S6). Classifiers’ performances varied sharply between participants with classification scores above 60% for some participants, while for other participant’s the classifier presented with chance-level performance.

**Figure 6:**
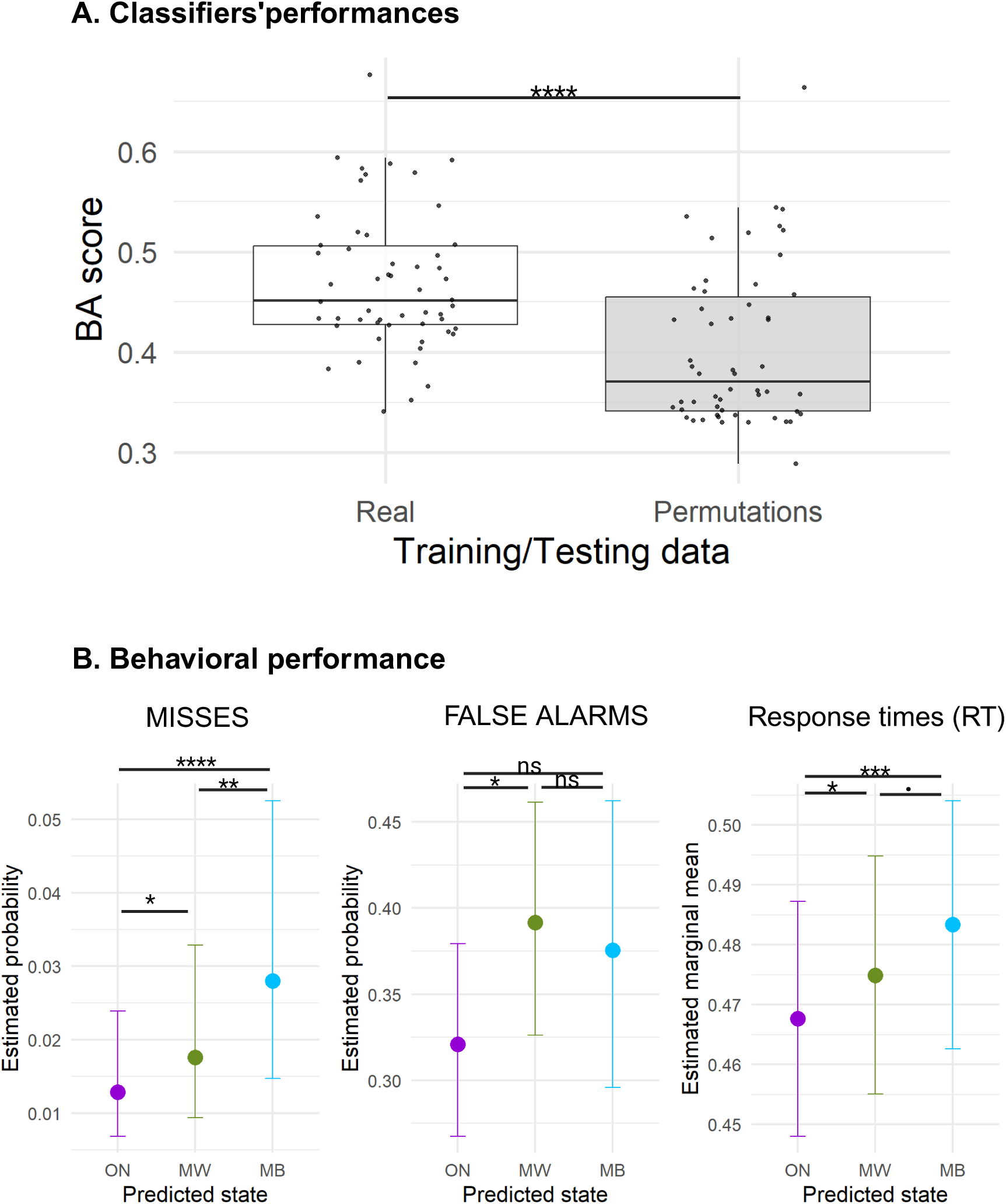
Trial-by-trial prediction of mind-state using a multivariate combination of neural markers. Panel A: Distribution of performances (balanced accuracy) of subject-level classifiers, both for true data and permuted data (mean score of 500 labels permutations). The classifier was trained and tested by cross-validation using labeled trials (<5 seconds before probe onsets). Spectral, complexity, connectivity, and ERP markers were used as raw features, followed by a dimensionality reduction by (non-linear) PCA. Only subjects presenting with the 3 mind-states were included in this analysis. Panel B: Behavioral markers as a function of the predicted mind state in non-labeled trials (>5s before probe onsets). (Binomial) linear mixed models were computed with predicted mind-state as the main explanatory factor, and subject ID/dataset as random intercepts. Statistical bars represent pairwise statistical comparisons between states. We found very similar behavioral patterns for predicted trials as compared to reported (labeled) trials. The detailed statistical results can be found in Table S6. ****: FDR corrected p-value<0.0001 ***: FDR corrected p-value<0.001 **: FDR corrected p-value<0.01 *: FDR corrected p-value<0.05 . : FDR corrected p-value<0.1 n.s.: FDR corrected p-value>0.1

We then used our trained classifiers to estimate the mind-state in non-labeled trials (>5s before probe onsets). To estimate the reliability of these single-trial predictions, we computed metrics of behavioral performance in these trials for each predicted state (O^--^N, M^--^W, M^--^B). Indeed, as shown previously, ON, MW and MB have different behavioral signatures so we can use behavior to check if these signatures are consistent with our predictions. And indeed, behavioral patterns of predicted states were very similar to the corresponding labeled states: we observed more misses in (predicted) M^--^B trials than in O^--^N and M^--^W trials, as well as longer RTs in M^--^B as compared to M^--^W and O^--^N trials (see Table S7 and Figure 6B). We also observed more FA in M^--^W compared to O^--^N trials but RT were significantly slower and not faster (see Figure 1). This pattern of results suggest that our classifiers were able to retrieve the behavioral signatures on ON, MW and MB states for unprobed trials.

## DISCUSSION

### Mind Blanking corresponds to a specific brain state, different from Mind Wandering

In this study replicating and extending a previously published study^7^, we examined the behavioral and neural correlates of MB, compared to those of MW and ON states. During a sustained attention task, we confirmed our previous findings^7^ showing that MB corresponds to a distinct mental state, characterized by a specific behavioral profile and by specific neural signatures. At a behavioral level, MB was characterized by response slowing and higher rate of misses, compatible with participants’ subjective report of having been “absent minded” during those moments. This behavioral profile was different from the one observed during MW that was characterized by an acceleration of responses and more false alarms, suggestive of an ‘impulsive’ pattern of behavior. These differences in behavioral outcomes between MW and MB were previously reported in different tasks (e.g., reading)^75^ or even task-free resting state (RT computed on probe responses)^60^. At the neural level, both state markers (spectral, complexity and connectivity measures) and brain responses to external stimuli differentiate MB from MW and ON states. This complements previous findings in fMRI associating spontaneous MB with a pattern of whole-brain hyper-connectivity and de-activation^21,23^. A multivariate combination of these neural metrics allowed for higher than chance prediction of mind-state on a trial-by-trial basis, further supporting the idea that MB, MW and ON states correspond to different mental and neurophysiological states. Taken together, these multimodal results reaffirm the specificity of MB as a distinct mental state^28^.

These results face the intrinsic limitations of the validity and reliability of self-reports during experience sampling^76^. First, nine participants never reported any instance of MB, raising the possibility that the occurrence of MB is trait-dependent, an idea supported by its higher prevalence in individuals with Attention-Deficit Hyperactivity Disorder (ADHD)^53,77,78^. It is also possible that individuals show different response biases in reporting MB. Second, since we used a probe-catching method where subjective reports were assessed every 1 min or so, we cannot exclude the possibility of mental contents/states happening during the inter-probe intervals that participants could not report. Third, the experience sampling method does not allow to determine precisely the time window that corresponds to a given mind-state. As a first step towards addressing this limitation in the future, we tried to predict the participant’s mind-state without reports using multivariate classifiers trained with neural metrics during those trials. Finally, our criterion for considering trials as belonging to a particular mind state (5 seconds) was motivated by technical reasons (see Methods), and not hypothesis or data-driven.

### Front vs Back dissociation between Mind Blanking and Mind Wandering

We found that spectral and complexity markers display opposite topographical profiles for MW and MB: whereas MB was characterized by an increase of both complexity and fast oscillatory activity over frontal electrodes (compared to MW), it was also characterized by a decrease in these same metrics over posterior electrodes. These results are in line with previous analyses in a subset of our dataset (40% of the current data)^7^, that found evidence for sleep-like slow-waves in both MW and MB, but with different regional localizations: while MW was associated with the presence of wake slow-waves over frontal scalp regions, MB was associated with their extension to posterior regions. Two previous fMRI studies showed that spontaneous MB is associated with an extensive cortical and thalamic de-activation as well as a widespread positive phase-coupling between brain regions^21,23^. Interestingly, chemogenetic inhibition of the mouse prefrontal cortex can result in (i) a decrease in neuronal firing, (ii) the presence of slow oscillations, (iii) an increase in functional connectivity^79^, which suggests that the different neural correlates of MB (slow waves, deactivation and hyper-connectivity) could reflect the same neural mechanisms.

These results contrast with another study in fMRI focusing this time on *voluntary* MB, which showed its association with a deactivation of lateral prefrontal regions and hippocampus, but an activation of anterior cingulate cortex^22^. While our spectral and complexity measures were computed in sensor space, and therefore lack precise anatomical information, the observed increase in fast oscillatory activity and neural complexity over centro-anterior electrodes during MB could also be related to ACC activation. These findings are also in line with studies contrasting episodes of MW in which participants are aware or unaware of their own MW. When caught unaware, participants exhibited more activation in frontal areas (dorsal Anterior Cingulate Cortex and ventral Pre Frontal Cortex)^80^, which could be related to the increase in fast oscillations we observed in MB, possibly suggesting a continuum between MB and unaware MW. Overall these results highlight the necessity to refine the phenomenological characterization of MB in order to better apprehend its neural correlates^18,19^. It further enriches the literature on spontaneous thoughts and experiences, which previously focused on MW but stressed the importance of considering phenomenal dimensions of MW such as meta-awareness, voluntariness or emotional valence. With MB, we emphasise that the continuous presence of experiential contents should not be taken for granted.

### Increased phase synchrony with decreased information sharing between distant brain regions during Mind Blanking

To date, previous studies have reported mixed results regarding functional connectivity during MB. A previous study using fMRI^22^ revealed lower functional connectivity between the default mode network and frontal, visual, and salience networks during voluntary MB as compared to MW. By contrast, using a coherence-based measure, a recent study^23^ found a pattern of global positive connectivity during spontaneous MB. Beyond potential differences between voluntary and spontaneous MB, it is important to consider that functional connectivity results depend very much on the type of measure being computed. In the present study, we computed two different metrics: the PLV^81^, a classical coherence-based measure that captures strictly linear correlations, and the wSMI, an information theory based measure which favors non-linear correlations over purely linear ones^45^. While the PLV showed an increase of inter-areal connectivity in MB as compared to both ON and MW (in line with the previously mentioned study), the wSMI revealed the inverse pattern, with reduced inter-areal (and more specifically, fronto-parietal) connectivity. Dissociations between coherence-based measures and the wSMI have already been reported, for example when contrasting Wakefulness and N3 sleep, where the whole-brain wSMI was significantly decreased in N3 sleep compared to wakefulness, whereas the wPLI (weighted Phase Lag Index) was not^82^. Our results could reflect a dissociation between increased phase-synchrony with decreased information sharing in MB, which could be caused by the occurrence of synchronous episodes of slow waves and neural silencing over associative cortices^83,84^. More frequent regional sleep-like slow waves in MB could explain the increase in PLV as these events would realign the phase of EEG signals across sources^7,85^. Several lines of evidence suggest that MB may mark the beginning of the transition into sleep. Sleep onset is not a discrete event but a gradual and multifaceted process^86^. Consistent with this view, MB is associated with behavioral slowing (increased misses and longer reaction times), alterations in brain connectivity and fronto-parietal sensory processing, and a shift toward sleep-like neural activity^7^. MB also increases with sleep deprivation^60^. However, it is important to emphasize that participants remained responsive to the SART constant visual stimulation, with fewer than 20% misses in the MB condition. Thus, if MB reflects a state closer to sleep, it still occurs within a globally wakeful state from both behavioral and physiological perspectives^87^. Lastly, this model stresses the influence of episodes of hypo-arousal in the occurrence of MB but does not exclude that MB could also occur in states of hyper-arousal as suggested elsewhere^60^.

### Disrupted neural representations of external stimuli during MB

Stimulus-induced activity analyses (ERPs, source reconstruction and temporal decoding) revealed similar patterns of processing of task stimuli during ON-task and MW states, significantly different from those observed during MB.

The most obvious difference between mind-states concerned the P3b component. A P3b response (central positivity between 400 and 600 ms post-stimulus) was clearly present during ON-task, was reduced but still observable during MW and seemed to lack during MB trials. While the neural correlates and signatures of the conscious processing of external information remain highly debated^88,89^, previous studies^47,70,71^ have proposed the P3b component as an EEG signature of conscious access. This interpretation remains debated, since some authors have claimed that the P3b relates more to post-perceptual processes associated with (external) report than to conscious awareness *per se*^90,91^. The reduction of the P3b observed in MW (shorter duration and decreased amplitude compared to ON-task) is in line with previous findings showing the impact of top-down attention on the P3b^92^. This reduction suggests the superposition of stimulus-unrelated processes during the time-window of the P3b response during MW. This last interpretation is in agreement with a recent study^69^, which demonstrated that the P3b can be modulated by participants’ attentional focus but conscious perception is always associated with a P3b-like activation of a broad set of associative cortical areas (frontal, parietal and temporal) even in the absence of report. When participants were paying attention and reported on the stimuli, additional cortical areas becoming involved during the same time window lead to a full-fledged P3b. Our source localization results align with this model with a similar activation of the dorsal and ventral visual streams during the late time-window for both ON and MW conditions, reaching some frontal areas in both conditions, but with a less spatially extended response during MW. Crucially, MB showed here a very different ERP signature at the sensor and source level. The relative absence of a P3b response as well as the absence of clear activation of the ventral/dorsal visual streams strongly suggests the absence of conscious access to external visual stimuli in MB trials in contrast with both ON and MW trials.

We also observed differences at the level of early visual processing between the different mind states. While no major differences were observable in sensor-space ERPs during the early time-window (<300ms post-stimulus), our temporal decoding analysis revealed the emergence of distinct visual neural representations around 150-200ms post-stimulus for both ON and MW trials (significant decoding against chance level performance of stimulus category), while this was not the case for MB trials (despite the fact that the number of trials were balanced across conditions). The absence of a significant encoding of visual stimuli over occipital areas around 200ms could suggest a disruption of the actualization of visual representations during MB. This aligns with the pattern of global cortical deactivations reported during MB^21^. It further suggests that the lack of late potentials observed during MB could be due to a weaker or absent sensory activation, failing to ignite the cascade of activations observed in ON and MW, albeit with decreased amplitude for the latter.

### Probing mind-state without external reports

The stream of conscious experiences is private and probing for its contents and dynamic is only possible with a sparse and disruptive sampling approach. We attempted here to bypass these reports by leveraging our new correlates of MW and MB. We showed first that we could predict above chance the mind-state category of trials just before a probe (using the mind-state reported following the probe as a ground truth) using a multivariate combination of different neural markers. Second, we applied the same algorithm to predict mind-states for trials away from the probes, so during moments where we do not have participant’s mind-state reports. We could retrieve the behavioral signatures of these states, which suggests that this MVPA approach partially captures a similar dynamics of conscious experience as evidenced in trials in which participants reported on their mental state. While the accuracy of our classifier approach remains very limited, this proof of concept provides an interesting new approach to estimate the second-by-second dynamics of mental states beyond the classical minute-by-minute sampling of subjective experience, paving the way for a fine-grained exploration of the dynamics of consciousness, without under-sampling or interfering with these dynamics by requiring a verbal report. Incorporating more detailed descriptions of mental states or embracing a multidimensional description of subjective experience could enhance both the accuracy and generalizability of such classifiers across individuals or groups, as recently shown in the context of fMRI studies^11,93^.

### Conclusion: is Mind Blanking a state without conscious content during wakefulness?

In the introduction of this paper, we presented the predicted neural correlates of a potential contentless conscious state during wakefulness, based on theoretical^34,35,94^ and empirical^31,39,45,46,69^ considerations. We showed here that the state of MB, a phenomenological, behavioral and neurophysiological distinct state from MW, fulfils these predictions. First, as predicted, the sharing of information between distant cortical areas was disrupted during MB, in particular in the delta and theta frequency bands. Second, the neural representations of external stimuli were significantly disrupted during MB, starting from early periods of processing and echoing into late processing, usually associated with conscious access, with a lack of the usual signatures of conscious access during this mind-state. Finally, spectral, complexity and coherence based connectivity metrics point towards the hypothesis of neural silencing of posterior associative cortices, a key node for consciousness according to most theoretical accounts^35,95^. Since we studied MB in a specific context and task-design, it is possible that some of the markers obtained here would not generalise to other instances of MB. Yet, some of the markers evidenced in this study are compatible with those obtained using a different neuroimaging technique (fMRI) and without a task^21,23^.

The presence or absence of internally generated representations during this mind-state remains an open question, and the impression of an “empty mind” reported by the participants during MB could be accounted for by different mechanisms (i.e., lack of metacognitive awareness, lack of memory encoding, language limitations) other than a lack of conscious experience altogether^19^. Still, based on the above presented results, we argue in favour of the more radical interpretation of MB as a “content-free” mind state. This would challenge our intuition of a continuous conscious content during wakefulness. In this view, conscious experience would be a discrete phenomenon, with discrete temporal islets with conscious content, separated by brief contentless periods^94^. While puzzling, this perspective would bring even more value to those precious moments of conscious experience.

## ONLINE METHODS

Sixty-eight healthy adults participated in a modified visual Sustained Attention to Response Task (SART), conducted over approximately 100 minutes, during which high-density EEG and oculometric data were continuously recorded. At various points during the task, participants were interrupted and asked to report their current mental state: specifically, whether they were focused on the task (task-focused), thinking about something unrelated to the task (mind-wandering), or not thinking about anything in particular (mind-blanking). We then compared behavioral and EEG data preceding these interruptions across the three reported mental states. A detailed account of the experimental procedures and data analyses is provided in the Supplementary Materials.

## Acknowledgments

AWC acknowledges the support of the Three Springs Foundation. The research leading to these results has received funding from the national program “Investissements d’avenir” ANR-10-IAIHU-0006. Part of this work was carried out in the CENIR of ICM. We gratefully acknowledge Laurent Hugueville and Amandine Carrie for their contribution to data collection at the Paris Brain Institute. We thank Teigane Mackay for data collection at Monash University, Naotsugu Tsuchiya, and Jennifer Windt for their support to TA at the inception of this line of research, and Jakob Howhy, Delphine Oudiette and Jacobo Sitt for the discussion about the nature and dynamics of consciousness. We thank the two anonymous reviewers for their constructive comments, as well as the late Prof. Jonathan Smallwood, who also reviewed this article. In addition to his thoughtful feedback on our work, we wish to honour Prof. Smallwood’s scientific contributions, which have deeply shaped this field and on which the present study directly builds.

## Author contributions

Conceptualization: EMM, AWC, LN, TA

Data acquisition: ALC, TA

Methodology: EMM, AWC, LB, LN, TA

Investigation: EMM, TA

Visualization: EMM, TA

Funding acquisition: LN, TA

Supervision: LN, TA

Writing – original draft: EMM

Writing – review & editing: EMM, ALC, AWC, LB, LN, TA

## Data and materials availability

All data, code, and materials used in the analysis are available here: 10.5281/zenodo.17649277.

## REFERENCES

1. James, W. The Principles of Psychology. (Henry Holt and Company, New York, 1890).

2. Smallwood, J., Obonsawin, M. & Heim, D. Task unrelated thought: the role of distributed processing. Conscious Cogn 12, 169–189 (2003).

3. Smallwood, J. & Schooler, J. W. The Science of Mind Wandering: Empirically Navigating the Stream of Consciousness. Annual Review of Psychology 66, 487–518 (2015).

4. Gruberger, M., Simon, E., Levkovitz, Y., Zangen, A. & Hendler, T. Towards a Neuroscience of Mind-Wandering. Frontiers in Human Neuroscience 5, (2011).

5. Compton, R. J., Gearinger, D. & Wild, H. The wandering mind oscillates: EEG alpha power is enhanced during moments of mind-wandering. Cogn Affect Behav Neurosci 19, 1184–1191 (2019).

6. Braboszcz, C. & Delorme, A. Lost in thoughts: neural markers of low alertness during mind wandering. Neuroimage 54, 3040–3047 (2011).

7. Andrillon, T., Burns, A., Mackay, T., Windt, J. & Tsuchiya, N. Predicting lapses of attention with sleep-like slow waves. Nat Commun 12, 3657 (2021).

8. Turnbull, A. et al. Left dorsolateral prefrontal cortex supports context-dependent prioritisation of off-task thought. Nature Communications 10, (2019).

9. Konu, D. et al. A role for the ventromedial prefrontal cortex in self-generated episodic social cognition. NeuroImage 218, 116977 (2020).

10. Sormaz, M. et al. Default mode network can support the level of detail in experience during active task states. Proceedings of the National Academy of Sciences 115, 9318–9323 (2018).

11. Wallace, R. S. et al. Mapping patterns of thought onto brain activity during movie-watching. eLife 13, RP97731 (2025).

12. Schooler, J. W. et al. Meta-awareness, perceptual decoupling and the wandering mind. Trends in Cognitive Sciences 15, 319–326 (2011).

13. Christoff, K., Irving, Z. C., Fox, K. C. R., Spreng, R. N. & Andrews-Hanna, J. R. Mind-wandering as spontaneous thought: a dynamic framework. Nature Reviews Neuroscience 17, 718–731 (2016).

14. Smallwood, J. et al. The neural correlates of ongoing conscious thought. iScience 24, 102132 (2021).

15. Ward, A. & Wegner, D. Mind-blanking: when the mind goes away. Frontiers in Psychology 4, (2013).

16. Schooler, J. W., Reichle, E. D. & Halpern, D. V. Zoning Out while Reading: Evidence for Dissociations between Experience and Metaconsciousness. in Thinking and seeing: Visual metacognition in adults and children 203–226 (MIT Press, Cambridge, MA, US, 2004).

17. Van Calster, L., D’Argembeau, A., Salmon, E., Peters, F. & Majerus, S. Fluctuations of Attentional Networks and Default Mode Network during the Resting State Reflect Variations in Cognitive States: Evidence from a Novel Resting-state Experience Sampling Method. Journal of Cognitive Neuroscience 29, 95–113 (2017).

18. Kaufmann, A., Parmigiani, S., Kawagoe, T., Zabaroff, E. & Wells, B. Two models of mind blanking. Eur J of Neuroscience ejn.16164 (2023) doi:10.1111/ejn.16164.

19. Fell, J. What is mind blanking: A conceptual clarification. Eur J of Neuroscience 56, 4837–4842 (2022).

20. Boulakis, P. A. & Demertzi, A. Relating mind-blanking to the content and dynamics of spontaneous thinking. Current Opinion in Behavioral Sciences 61, 101481 (2025).

21. Boulakis, P. A., Mortaheb, S., Van Calster, L., Majerus, S. & Demertzi, A. Whole-Brain Deactivations Precede Uninduced Mind-Blanking Reports. J. Neurosci. 43, 6807–6815 (2023).

22. Kawagoe, T., Onoda, K. & Yamaguchi, S. The neural correlates of “mind blanking”: When the mind goes away. Human Brain Mapping 40, 4934–4940 (2019).

23. Mortaheb, S. et al. Mind blanking is a distinct mental state linked to a recurrent brain profile of globally positive connectivity during ongoing mentation. Proceedings of the National Academy of Sciences 119, e2200511119 (2022).

24. Mittner, M., Hawkins, G. E., Boekel, W. & Forstmann, B. U. A Neural Model of Mind Wandering. Trends in Cognitive Sciences 20, 570–578 (2016).

25. Unsworth, N. & Robison, M. K. Tracking arousal state and mind wandering with pupillometry. Cogn Affect Behav Neurosci 18, 638–664 (2018).

26. Andrillon, T. et al. Does the Mind Wander When the Brain Takes a Break? Local Sleep in Wakefulness, Attentional Lapses and Mind-Wandering. Frontiers in Neuroscience 13, (2019).

27. Andrillon, T., Lutz, A., Windt, J. & Demertzi, A. Where is my Mind?: A Neurocognitive Investigation of Mind Blanking. Preprint at 10.31234/osf.io/xmtga (2024).

28. Boulakis, P. A. & Demertzi, A. Relating mind-blanking to the content and dynamics of spontaneous thinking. Current Opinion in Behavioral Sciences 61, (2025).

29. Siclari, F., LaRocque, J., Postle, B. & Tononi, G. Assessing sleep consciousness within subjects using a serial awakening paradigm. Frontiers in Psychology 4, (2013).

30. Siclari, F. et al. The neural correlates of dreaming. Nature Neuroscience 20, 872–878 (2017).

31. Türker, B. et al. Behavioral and brain responses to verbal stimuli reveal transient periods of cognitive integration of the external world during sleep. Nat Neurosci 26, 1981–1993 (2023).

32. Mazza, S. et al. Asleep but aware? Brain and Cognition 87, 7–15 (2014).

33. Andrillon, T. & Kouider, S. The vigilant sleeper: neural mechanisms of sensory (de)coupling during sleep. Current Opinion in Physiology (2019) doi:10.1016/j.cophys.2019.12.002.

34. Dehaene, S. Towards a cognitive neuroscience of consciousness: basic evidence and a workspace framework. Cognition 79, 1–37 (2001).

35. Mashour, G. A., Roelfsema, P., Changeux, J.-P. & Dehaene, S. Conscious Processing and the Global Neuronal Workspace Hypothesis. Neuron 105, 776–798 (2020).

36. Tononi, G. & Massimini, M. Why does consciousness fade in early sleep? Ann N Y Acad Sci 1129, 330–334 (2008).

37. Casali, A. G. et al. A Theoretically Based Index of Consciousness Independent of Sensory Processing and Behavior. Science Translational Medicine 5, 198ra105–198ra105 (2013).

38. Casarotto, S. et al. Exploring the Neurophysiological Correlates of Loss and Recovery of Consciousness: Perturbational Complexity. in Brain Function and Responsiveness in Disorders of Consciousness (eds. Monti, M. M. & Sannita, W. G.) 93–104 (Springer International Publishing, Cham, 2016). doi:10.1007/978-3-319-21425-2_8.

39. Sitt, J. D. et al. Large scale screening of neural signatures of consciousness in patients in a vegetative or minimally conscious state. Brain 137, 2258–2270 (2014).

40. Engemann, D. A. et al. Robust EEG-based cross-site and cross-protocol classification of states of consciousness. Brain 141, 3179–3192 (2018).

41. Boveroux, P. et al. Breakdown of within- and between-network resting state functional magnetic resonance imaging connectivity during propofol-induced loss of consciousness. Anesthesiology 113, 1038–1053 (2010).

42. Ranft, A. et al. Neural Correlates of Sevoflurane-induced Unconsciousness Identified by Simultaneous Functional Magnetic Resonance Imaging and Electroencephalography. Anesthesiology 125, 861–872 (2016).

43. Horovitz, S. G. et al. Decoupling of the brain’s default mode network during deep sleep. Proc Natl Acad Sci U S A 106, 11376–11381 (2009).

44. Spoormaker, V. I. et al. Development of a large-scale functional brain network during human non-rapid eye movement sleep. J Neurosci 30, 11379–11387 (2010).

45. King, J.-R. et al. Information Sharing in the Brain Indexes Consciousness in Noncommunicative Patients. Current Biology 23, 1914–1919 (2013).

46. Bourdillon, P. et al. Brain-scale cortico-cortical functional connectivity in the delta-theta band is a robust signature of conscious states: an intracranial and scalp EEG study. Scientific Reports 10, 1–13 (2020).

47. Bekinschtein, T. A. et al. Neural signature of the conscious processing of auditory regularities. PNAS 106, 1672–1677 (2009).

48. Perez, P. et al. Auditory Event-Related “Global Effect” Predicts Recovery of Overt Consciousness. Frontiers in Neurology 11, (2021).

49. King, J.-R., Gramfort, A., Schurger, A., Naccache, L. & Dehaene, S. Two Distinct Dynamic Modes Subtend the Detection of Unexpected Sounds. PLoS ONE 9, e85791 (2014).

50. Strauss, M. et al. Disruption of hierarchical predictive coding during sleep. Proceedings of the National Academy of Sciences 112, E1353–E1362 (2015).

51. Hayat, H. et al. Reduced neural feedback signaling despite robust neuron and gamma auditory responses during human sleep. Nat Neurosci 25, 935–943 (2022).

52. Pérez, P. et al. Content–state dimensions characterize different types of neuronal markers of consciousness. Neuroscience of Consciousness 2024, niae027 (2024).

53. Van den Driessche, C., et al. Attentional Lapses in Attention-Deficit/Hyperactivity Disorder: Blank Rather Than Wandering Thoughts. Psychol Sci 28, 1375–1386 (2017).

54. Richman, J. S. & Moorman, J. R. Physiological time-series analysis using approximate entropy and sample entropy. American Journal of Physiology-Heart and Circulatory Physiology 278, H2039–H2049 (2000).

55. Engemann, D. A. et al. Robust EEG-based cross-site and cross-protocol classification of states of consciousness. Brain 141, 3179–3192 (2018).

56. Schartner, M. M. et al. Global and local complexity of intracranial EEG decreases during NREM sleep. Neuroscience of Consciousness 2017, (2017).

57. Türker, B. et al. Behavioral and brain responses to verbal stimuli reveal transient periods of cognitive integration of external world in all sleep stages. 2022.05.04.490484 Preprint at 10.1101/2022.05.04.490484 (2022).

58. Munoz Musat, E., Rohaut, B., Sangare, A., Benhaiem, J.-M. & Naccache, L. Hypnotic Induction of Deafness to Elementary Sounds: An Electroencephalography Case-Study and a Proposed Cognitive and Neural Scenario. Frontiers in Neuroscience 16, (2022).

59. Martínez Vivot, R., Pallavicini, C., Zamberlan, F., Vigo, D. & Tagliazucchi, E. Meditation Increases the Entropy of Brain Oscillatory Activity. Neuroscience 431, 40–51 (2020).

60. Boulakis, P. A. et al. Variations of autonomic arousal mediate the reportability of mind blanking occurrences. Sci Rep 15, 4956 (2025).

61. Bastuji, H. & García-Larrea, L. Evoked potentials as a tool for the investigation of human sleep. Sleep Med Rev 3, 23–45 (1999).

62. Ruby, P., Caclin, A., Boulet, S., Delpuech, C. & Morlet, D. Odd sound processing in the sleeping brain. J Cogn Neurosci 20, 296–311 (2008).

63. Nir, Y., Vyazovskiy, V. V., Cirelli, C., Banks, M. I. & Tononi, G. Auditory responses and stimulus-specific adaptation in rat auditory cortex are preserved across NREM and REM sleep. Cereb Cortex 25, 1362–1378 (2015).

64. Kouider, S., Andrillon, T., Barbosa, L. S., Goupil, L. & Bekinschtein, T. A. Inducing Task-Relevant Responses to Speech in the Sleeping Brain. Current Biology 24, 2208–2214 (2014).

65. Andrillon, T., Poulsen, A. T., Hansen, L. K., Léger, D. & Kouider, S. Neural Markers of Responsiveness to the Environment in Human Sleep. J. Neurosci. 36, 6583–6596 (2016).

66. Fischer, C. et al. Mismatch negativity and late auditory evoked potentials in comatose patients. Clinical Neurophysiology 110, 1601–1610 (1999).

67. Fischer, C., Luauté, J., Adeleine, P. & Morlet, D. Predictive value of sensory and cognitive evoked potentials for awakening from coma. Neurology 63, 669–673 (2004).

68. Robinson, L. R., Micklesen, P. J., Tirschwell, D. L. & Lew, H. L. Predictive value of somatosensory evoked potentials for awakening from coma*. Critical Care Medicine 31, 960 (2003).

69. Sergent, C. et al. Bifurcation in brain dynamics reveals a signature of conscious processing independent of report. Nat Commun 12, 1149 (2021).

70. Sergent, C., Baillet, S. & Dehaene, S. Timing of the brain events underlying access to consciousness during the attentional blink. Nature Neuroscience 8, 1391–1400 (2005).

71. Del Cul, A., Baillet, S. & Dehaene, S. Brain Dynamics Underlying the Nonlinear Threshold for Access to Consciousness. PLoS Biol 5, (2007).

72. Gaillard, R. et al. Converging Intracranial Markers of Conscious Access. PLoS Biol 7, (2009).

73. Spehlmann, R. The averaged electrical responses to diffuse and to patterned light in the human. Electroencephalogr Clin Neurophysiol 19, 560–569 (1965).

74. King, J.-R., et al. Encoding and Decoding Neuronal Dynamics: Methodological Framework to Uncover the Algorithms of Cognition. (2018).

75. Ward, A. F. & Wegner, D. M. Mind-blanking: when the mind goes away. Front Psychol 4, 650 (2013).

76. Nisbett, R. E. & Wilson, T. D. Telling more than we can know: Verbal reports on mental processes. Psychological Review 84, 231–259 (1977).

77. Pinggal, E. et al. Sleep-like Slow Waves During Wakefulness Mediate Attention and Vigilance Difficulties in Adult Attention-Deficit/Hyperactivity Disorder. Preprint at 10.1101/2025.07.27.666103 (2025).

78. Madiouni, C., Lopez, R., Gély-Nargeot, M.-C., Lebrun, C. & Bayard, S. Mind-wandering and sleepiness in adults with attention-deficit/hyperactivity disorder. Psychiatry Research 287, 112901 (2020).

79. Rocchi, F. et al. Increased fMRI connectivity upon chemogenetic inhibition of the mouse prefrontal cortex. Nat Commun 13, 1056 (2022).

80. Christoff, K., Gordon, A. M., Smallwood, J., Smith, R. & Schooler, J. W. Experience sampling during fMRI reveals default network and executive system contributions to mind wandering. Proceedings of the National Academy of Sciences of the United States of America 106, 8719–8724 (2009).

81. Aydore, S., Pantazis, D. & Leahy, R. M. A note on the phase locking value and its properties. NeuroImage 74, 231–244 (2013).

82. Imperatori, L. S. et al. EEG functional connectivity metrics wPLI and wSMI account for distinct types of brain functional interactions. Sci Rep 9, 8894 (2019).

83. Aedo-Jury, F., Schwalm, M., Hamzehpour, L. & Stroh, A. Brain states govern the spatio-temporal dynamics of resting-state functional connectivity. eLife 9, e53186 (2020).

84. El-Baba, M. et al. Functional connectivity dynamics slow with descent from wakefulness to sleep. PLOS ONE 14, e0224669 (2019).

85. Pigorini, A. et al. Bistability breaks-off deterministic responses to intracortical stimulation during non-REM sleep. NeuroImage 112, 105–113 (2015).

86. Lacaux, C., Strauss, M., Bekinschtein, T. A. & Oudiette, D. Embracing sleep-onset complexity. Trends in Neurosciences 47, 273–288 (2024).

87. Andrillon, T. & Oudiette, D. What is sleep exactly? Global and local modulations of sleep oscillations all around the clock. Neuroscience & Biobehavioral Reviews 155, 105465 (2023).

88. Aru, J., Bachmann, T., Singer, W. & Melloni, L. Distilling the neural correlates of consciousness. Neuroscience & Biobehavioral Reviews 36, 737–746 (2012).

89. Tsuchiya, N., Wilke, M., Frässle, S. & Lamme, V. A. F. No-Report Paradigms: Extracting the True Neural Correlates of Consciousness. Trends in Cognitive Sciences 19, 757–770 (2015).

90. Cohen, M. A., Ortego, K., Kyroudis, A. & Pitts, M. Distinguishing the Neural Correlates of Perceptual Awareness and Postperceptual Processing. J. Neurosci. 40, 4925–4935 (2020).

91. Pitts, M. A., Metzler, S. & Hillyard, S. A. Isolating neural correlates of conscious perception from neural correlates of reporting one’s perception. Front Psychol 5, 1078 (2014).

92. Polich, J. Updating P300: An integrative theory of P3a and P3b. Clinical Neurophysiology 118, 2128–2148 (2007).

93. Mckeown, B. et al. Self-reports map the landscape of task states derived from brain imaging. Commun Psychol 3, 8 (2025).

94. Naccache, L. Why and how access consciousness can account for phenomenal consciousness. Philosophical Transactions of the Royal Society B: Biological Sciences 373, 20170357 (2018).

95. Tononi, G. The Integrated Information Theory of Consciousness: An Updated Account. Archives Italiennes de Biologie 150, 56–90 (2012).

