## Supplementary Figures and Table for "Behavioral, experiential, and physiological signatures of mind blanking"

### SUPPLEMENTARY METHODS

#### Experimental procedure and data collection

##### *Participants*

A total of sixty-eight (N = 68) healthy adult subjects were enrolled in this study. Thirty-two (N=32) were recorded at the Monash University (Melbourne, Australia), between June 2018 and February 2019. An additional set of thirty-six (N=36) subjects were recorded at the Paris Brain Institute (Paris, France) from June 2024 to September 2024. The original experiment (Monash) was conducted in English, and instructions were translated in French for the Paris experiment. All other materials and procedures (including the EEG equipment) were similar for the two groups. Due to technical complications during the recording process or suboptimal physiological recording quality identified via retrospective visual examination of the data, six individuals were excluded from subsequent analysis. The final study group, comprising 62 participants (age: 27.9 ± 0.7 years, mean ± standard deviation (age missing for one individual); 31 females), was included in all analyses. Prior to study involvement, participants submitted written informed consent. Both sites Human Research Ethics Committees approved the study protocol (Monash University Human Research Ethics Committee, Project ID: 10994 ; and Comité d'Éthique de la Recherche (CER) de Sorbonne Université, Project ID: CER-2023-DEGRAVE-MW_Respi)

##### *Experimental design and stimuli*

Participants were positioned in a dimly lit room, seated approximately 60 cm away from a computer screen with their chin supported. Task instructions, stimuli presentation, and button response collection were executed using the Psychtoolbox extension for Matlab (Mathworks, Natick, MA, USA).

The experimental design involved two adapted SARTs, where participants were instructed to focus on a series of human face images during Face SART blocks and on digits during Digit SART blocks. The sequence of Face and Digit blocks was pseudo-randomized for each participant. Each block lasted for roughly 12-15 minutes, with rest periods permitted between blocks. Participants completed three Face SART and three Digit SART blocks, totalling a duration of 86.7 ± 1.2 minutes (mean ± standard deviation) for the SART, and 108.4 ± 3.4 minutes for the entire recording session. Prior to each type of Face and Digit SART, participants underwent a brief training session (27 trials) with performance feedback on correct trials and average reaction times (RTs). Participants were urged to prioritize accuracy over speed.

Face and digit stimuli were displayed continuously, with each stimulus shown for 750-1250 ms (uniform random jitter). Face stimuli were derived from the Radboud Face Database(1), consisting of eight neutral faces (four females, four males) and one smiling female face. Digits ranging from 1 to 9 were displayed in a fixed font size. For Face SART, participants pressed a button for all neutral faces (Go trials) and refrained from pressing the button for the smiling face (No Go trials). The face order was pseudo-randomized throughout the task (presentation order permuted every nine stimuli, without immediate repetition). For Digit SART, participants pressed a button for all digits except digit 3 (No Go trials), with the digit order pseudo-randomized similarly.

During the SART, stimulus presentation was halted at random intervals (every 40-70 s, uniform random jitter) accompanied by a sound and the word "STOP" on the screen. These interruptions allowed for the assessment of participants' mental states using a series of eight questions. Participants were notably instructed to report their attentional focus "just before the interruption" by selecting one of four options: (1) "task-focused" (ON), (2) "off-task" (mind-wandering, MW), (3) "mind-blanking" (MB), or (4) "don't remember." Since the fourth option accounted for only 2.6% of all probes (less than 2 probes per participant on average) and previous studies do not consistently distinguish between these options (2), we combined the third and fourth options as MB in all analyses. Whenever participants chose the off-task option, they were prompted to indicate the origin of their thoughts or experiences: (1) the environment (distractions), (2) the task (interferences), or (3) internally generated contents. Participants also rated their vigilance levels over the past few trials on a 4-point scale (1 = Extremely Sleepy, 4 = Extremely Alert). Each 12-15 minute SART block contained ten interruptions (30 interruptions per SART task, 60 interruptions per participant). Participants were briefed on the interruptions and question types prior to the experiment. The mental state categories (ON, MW, and MB) were explained both orally and in writing to participants.

##### *Physiological recordings*

High-density scalp EEG data were obtained utilizing an EasyCap (Monash) or actiCap Snap (Paris) with 63 active electrodes, connected to a BrainAmp system (Brain Products GmbH). A ground electrode was positioned frontally (Fpz in the 10-20 system), while electrodes were referenced online to a frontal electrode (AFz). Additional electrodes were placed above and below the left and right canthi to record ocular movements (electrooculogram, EOG). Two electrodes were placed over the deltoid muscles to capture electrocardiographic (ECG) activity. EEG, EOG, and ECG signals were sampled at a rate of 500 Hz. Eye movements and pupil size from one eye were recorded using an EyeLink 1000 system (SR Research) but not analyzed here. In the Paris dataset, participants were also equipped with a respiratory belt.

#### Behavioral analyses

Go trials were deemed incorrect (Miss) if no response was registered between the onset of the stimulus and the subsequent stimulus onset. In contrast, No-Go trials were classified as incorrect (false alarm; FA) if a response was detected within the same time frame. Reaction times (RTs) were calculated from the onset of stimulus presentation. Behavioral performances for each probe were computed using trials within 5 seconds from probe onset. The choice of a 5 second window allowed the inclusion of sufficient No-Go trials for analysis (in average one No-Go trial every two probes), while focusing on trials very close from probe onset and subsequent subjective reports.

#### EEG preprocessing and analysis

##### *Preprocessing*

The raw EEG signal was first high-pass filtered >1 Hz, using a linear-phase, zero-phase FIR (finite impulse response) filter to remove low-frequency drifts which could negatively affect the quality of the ICA fit (see below). A notch filter was then applied (stopbands: [49, 51], [99,101] Hz, FIR filter) to remove line noise. Electrodes that were visually identified as noisy throughout the recording were removed and interpolated using neighbouring electrodes. The EEG signal was then decomposed using independent component analysis (ICA), and the components containing ocular artifacts or electrocardiogram (ECG) artifacts were removed prior to signal reconstruction.

The ICA-cleaned EEG signal was then down-sampled to 250 Hz, and low-pass filtered <60 Hz. An average reference was applied to all EEG electrodes. Finally, the continuous EEG data were segmented in two different manners, so as to obtain two sets of epochs for different analyses: from -5 to 0 seconds relative to the probe onsets ("Probe epochs"), and from -0.25 to 0.75 seconds relative to the stimuli onsets ("Target epochs"). Probe epochs were labelled according to the mind state reported by the participant at probe onset. Target epochs in the 5 second period prior to the probes were labelled according to the subjective report, all other Target epochs were considered non-labelled (unknown mind state).

##### *Sensor space analyses*

*Calculation of electroencephalographic markers tracking cognitive fluctuations*

Previous work has shown that cognitive and consciousness state modifications can be tracked using different spectral, complexity or connectivity measures derived from the scalp or intracranial electroencephalographic recording. By combining these markers, it is possible to distinguish conscious participants, patients in a minimal consciousness state and patients with unresponsive wakefulness syndrome(3–5). These measures can also differentiate sleep stages(6–8) and track cognitive and consciousness modifications related to psychedelics or meditation(9). In our study, we computed these different measures to try to describe the neural correlates of the different mind states (ON, MW and MB). We therefore computed these measures on a trial-by-trial basis on “Probe epochs” (from -5s to 0s relative to probe onsets), the 5 second time-window allowing for both a sufficient time-period for correct maker computation and for (very) close proximity to the subjective reports.

Prior to marker computation, we performed a spatial Laplacian transformation of the EEG signal (also known as Current Source Density (CSD) estimate)(10). We applied a spherical spline surface Laplacian transformation, with 50 Legendre terms, a stiffness of the spline of value 4 and a regularization parameter (lambda 2) of 1e-5.

Spectral markers

We computed the normalized power spectral densities (PSD) in delta (1-4 Hz), theta (4-8 Hz), alpha (8-12 Hz), beta (12-30 Hz) and gamma (30-60) frequency bands, using Welch’s method. The length of each Welch segment (windowed with a Hamming window) was set to be equal to the length of the FFT (fast Fourier transform) and equal to 256 samples (1024 ms). The obtained segments were then averaged, in order to obtain a single value per epoch, channel and frequency. To obtain the normalized PSDs in each frequency band of interest, we: 1) added the raw power of all the frequencies in each frequency band of interest; and 2) computed for each trial and each electrode the normalized PSD by normalizing the raw frequency band PSD by the total power of the electrode.

Complexity markers

We computed two different complexity measures: the Kolmogorov Complexity (KC) and the Sample Entropy (SE). Although they differ in their method of calculation, both measure the level of predictability and regularity of the EEG signal: the more the signal is unpredictable, the more it’s algorithmic complexity is important (more information is needed to describe the signal), and the more the complexity of the local neural processing captured by the EEG sensor is assumed to be important.

We computed these two complexity measures for each trial and each EEG electrode (n= 64). See Supplementary material of Sitt et. al (2014)(4) for a detailed description of KC, as well as its computation. Details regarding the SE measure can be found in Richman & Moorman (2000)(11).

Connectivity markers

Weighted symbolic mutual information

We focused on the *weighted symbolic mutual information* (wSMI), a functional connectivity measure capturing linear and non-linear coupling between sensors, which relies on the symbolic transformation of the EEG signal. The wSMI assesses the degree to which two EEG sensors exhibit non-random joint fluctuations, implying information sharing. The time series X and Y in all EEG channels are initially converted into discrete symbol sequences (X^, Y^). These symbols are encoded based on the trends in amplitudes in a predetermined number of consecutive time points. The symbols are therefore defined by two parameters: the number of time points considered (k) and the temporal separation between the time points that constitute a symbol (**$\tau$**). Following previous work, we chose the parameter k to be 3, implying that the symbols are constituted of three elements, leading to 3! = 6 different potential symbols in total. The parameter **$\tau$** sets the broad frequency-specific window of sensitivity of this connectivity measure, allowing to compute the wSMI in the delta (**$\tau$**= 64ms), theta (**$\tau$**= 32ms) and alpha (**$\tau$**= 16ms) bands. For each frequency band, the joint probability of each pair of symbols co-occurring in two different time series is computed to estimate the *symbolic mutual information* (SMI) shared across two signals. To minimise volume conduction, the *weighted* symbolic mutual information disregards co-occurrences of identical or opposite-sign signals.

The wSMI between two EEG sensors is computed, after symbolic transformation, according to the following formula:

$$\mathrm{wSMI}(\widehat{X}, \widehat{Y}) = \frac{1}{log(k!)} \sum_{\widehat{x} \epsilon \widehat{X}} \sum_{\hat{y} \epsilon \widehat{Y}} w(\widehat{x}, \hat{y}) p(\widehat{x}, \hat{y}) log \frac{p(\widehat{x}, \hat{y})}{p(\widehat{x}) p(\hat{y})}$$

where w(x,y) is the weight matrix and p(x,y) is the joint probability of co-occurrence of symbol x in signal X and symbol y in signal Y. Finally p(x) and p(y) are the probabilities of those symbols in each signal.

We computed, in each frequency band, and for each trial, the wSMI between each pair of electrodes (N pairs=2016). To obtain the topographical maps shown in Supplementary Figure S1, we computed, for each sensor, its median connectivity with all other sensors, obtaining a single value per channel per trial.

Phase Locking Value (PLV)

We also computed a more classical spectral connectivity measure, the Phase Locking Value (PLV), on a trial-by-trial basis. For each channel $i$, we extracted the instantaneous phase $\phi i^{a}(t)$ of the analytical signal $xi^{a}(t)$ of the time series $xi\ (t)$. Then, for each pair $(i,j)$ of channels, we computed the modulus of the time averaged phase difference mapped onto the unit circle:

$$PLV_{ij} = \left| \frac{1}{T} \sum_{t} e^{i(\phi_i^a(t) - \phi_j^a(t))} \right|$$

As with wSMI, we computed the PLV at trial level for each pair of electrodes, and then, for topographical representation, we computed the median PLV per EEG channel per trial.

*Event-related potentials (ERPs) elicited by visual stimuli during the task*

In order to study the neural fate of visual stimuli (digits and faces) during the SART task as a function of mind state, we conducted an event-related potential analysis on “Target epochs” (-0.25 to 0.75 s relative to stimulus onset). We first applied a baseline correction relative to the -0.25 to 0s time interval. Given the unbalanced nature of our dataset (important variability of the distribution of mind states across participants), we performed a single-trial multi-level analysis of ERPs responses(12) (see Statistics section for more details about this analysis). Before statistical analysis, to reduce inter-subject variability and to better fit the statistical model, we normalized channel voltages independently for each subject, by subtracting to each point of the *channel x time* matrix the mean *m* and dividing the result by the standard deviation *s*, with *m* and *s* being computed across all points of the *channel x time* matrix. Given that trials follow one another in close succession, it is normal for the baseline activity to be influenced by the activity of the previous trials as seen in Figure 4.

As a control, we also conducted the previously described analysis for each stimulus type (Faces vs. Digits) separately (see Figure S3).

*Temporal decoding of stimulus type*

To further study the neural fate of the visual stimuli as a function of mind state, we assessed whether we could decode stimulus type (Faces vs. Digits) from brain responses using multivariate pattern analysis (MVPA), with temporal decoding methods(13–15). Briefly, this type of analysis consists in using a subset of the data to train a linear classifier at each time-point to differentiate trials where a *Face* was presented from trials where a *Digit* was presented, and then testing its performance, independently at each time-point, in a different subset of the data. This method provides statistics tracking the temporal dynamics of mental representations. In the present study, we were interested in the differential profile of the temporal dynamics of mental representations related to external stimuli, as a function of the subject's mind-states (ON,MW and MB).

First, to avoid potential biases related to an unbalanced number of trials between mind-states (example: significantly more trials belonging to the ON mind state would induce better classifier performance in this condition than in the MB condition, independently of differential neural stimulus processing between conditions), we randomly downsampled, for each subject, the number of trials, so each mind state was equally represented. Then, independently for each reported mind-state (ON, MW, MB), we standardized the data at the subject level to suppress interindividual variability. At each time sample, and independently for each channel, channel voltages were standardized across trials sampled from the same participant, by removing the mean and scaling to unit variance. We then trained a linear classifier to decode Digits vs. Faces, using an L2-regularized (C=1) logistic regression, in a 10-fold cross-validation procedure. In each fold, all the trials were shuffled in a pseudo-randomized manner and split into a training set (9/10 of the trials) and a testing set (1/10 of the trials). The classifier's features were the standardized channel amplitudes. This training procedure was applied at each time step independently. Following the temporal decoding approach, the model trained at each time step was then tested at the same time-step on the testing set trials, at each cross-validation fold. The classifier performance at each testing time was evaluated by the area under the receiver operating curve (AUC) at each cross-validation fold. Global performance for each condition and at each time point was obtained by averaging the AUCs across folds. At each time point, performance against chance level was computed (see Statistics section). For visualization purposes, performance curves in function of time were smoothed using a gaussian filter, this filter wasn't applied for statistical analysis.

*Decoding of mind-state using dimensionality reduction and a Random Forest classifier*

Training and testing the classifier on labeled trials

We set out to determine if the participant's mind state could be predicted based on brain activity, on a trial-by-trial basis, using machine learning classifiers. We first focused on "Labeled Target" epochs, where the participant's mind state was considered as known (trials in the 5 seconds before subjective reports). From these epochs, we extracted the previously described EEG metrics (spectral, complexity, and connectivity, see "Calculation of electroencephalographic markers tracking cognitive modifications" section). We also added two supplementary features, the mean channel voltages during the early time-period (100 ms to 300ms relative to trial onset) and the mean channel voltages during the late time-period (400-600ms relative to trial onset). For all the previously mentioned markers, we extracted for each trial the values of these markers for each EEG channel (obtaining thus 14 x 64 = 896 features). Given the very high dimensionality of our feature space, we implemented a dimensionality reduction technique prior to classification, using a non-linear (kernel) Principal Component Analysis (PCA)(16). Classification was performed afterwards using Random Forest Classifiers. To take into account individual variability in neural metrics, we trained and tested a PCA/Random Forest classifier independently per participant, the hyperparameters of the PCA (Kernel type, number of components) and of the Random Forest (number of estimators, criterion, maximum depth) being fitted independently for each participant (see below). Importantly, to ensure an appropriate number of components/number of features ratio, the number of PCA components was chosen in the interval [2,25], corresponding therefore to a ratio of 1 component/10 trials or less, and hence avoiding overfitting. Kernel type was chosen from a list including the following Kernels: Linear, Polynomial, RBF and Sigmoid.

To find the best hyperparameters for our PCA/Random Forest pipeline, while also avoiding overfitting, we adopted a "Randomized Grid Search/Cross-validation procedure", using 100 iterations. We performed this procedure for each participant. In each iteration, a combination of parameter settings was randomly sampled from all specified possible values of each parameter, then the PCA/Random Forest classifier was trained and tested by cross-validation (see below). At the end of all iterations, the set of hyperparameters that obtained the best cross-validated score (balanced accuracy, see below) was selected for further analyses. Comparison to chance level performance was obtained using a 500-permutation procedure, independently for each participant. First, the true classifier's score was obtained by training and testing the best model (best set of hyperparameters) by cross validation. Then, at each permutation, trial labels were randomly shuffled, and the entire cross-validation procedure was repeated. We thus obtained a distribution of chance-level scores, for each participant. For each participant, we estimated the individual significance against chance by computing a p-value (by counting the number of permutation scores equal or higher to the true score and dividing it by the number of permutations plus one). Group-level significance against chance was also computed, by comparing the distribution of the classifiers' true scores across participants to the distribution of the mean permutation scores across participants (see Statistics section).

Cross-validation was performed using a 5-stratified group fold strategy. In each fold, 4/5 segments of the data were used to train the classifier and the remaining segment was used to test the classifier (computation of the balanced accuracy score, see below). Note that class distributions (proportion of ON, MW, and MB trials) were preserved across partitions of the data. Note also that folds were grouped by experimental block (6 block per participant): the blocks that served for the training were not used for the testing. This was done to avoid overfitting linked to temporal proximity of epochs (trials) situated in the same experimental blocks. Note also that this group parameter (experimental block) was also used during the permutation procedure (labels were shuffled inside each block), to avoid over-estimation of significance against chance. At the end of the cross-validation procedure, mean performance scores were obtained by averaging the values obtained in each fold.

We computed a performance score adapted to unbalanced datasets such as ours (unbalanced number of ON, MW, and MB trials): the balanced accuracy score. The balanced accuracy is defined, in multi-class classification, as the average sensitivity Se ("How many relevant items are retrieved?") for each class. The sensitivity is computed as the number of true positives, divided by the number of true positives plus false negatives. Hence, the balanced accuracy (BA) was computed in our dataset according to the following formula:

$$_{BA} = \frac{Se_{ON} + Se_{MW} + Se_{MB}}{3} = \frac{(TP/TP+FN)_{ON} + (TP/TP+FN)_{MW} + (TP/TP+FN)_{MB}}{3}$$

Note again that all the previously presented steps were performed using the Labeled Target epochs, where the participant's mind state was considered as known (trials in the 5 second period before subjective reports).

We also computed other very used classification metrics, at subject level and for each class: the Se (also called recall), the precision (TP/TP+FP) and the F1-score ((precision x recall)/(precision + recall)).

Predicting behavioral performances in non-labeled trials

To further test the performances of our classifier, trained on labeled trials, we predicted the mind state in non-labeled trials (more than 5 seconds before probe onsets). Practicaly speaking, we first fitted individual models with labeled data, and then predicted the mind state in non-labeled trials. Given the very high number of non-labeled trials (n=182,970) and its associated computational cost, we randomly subsampled, independently for each subject, 10% of non-labeled trials, resulting in a total of 18,293 trials included in this analysis. To assess the accuracy of this prediction, we once again computed behavioral performances, but this time only on non-labeled trials, and compared the behavioral profile as a function of the mind-state in non-labeled trials (according to the classifier's prediction), to the behavioral profile in labeled trials.

##### *Source space analyses*

*Forward Model and Source Modeling*

We used a constrained distributed model consisting of 15,000 current dipoles, constrained to the cortical mantle of a generic brain model obtained from the BrainStorm software package. EEG forward modeling was computed using a symmetric Boundary Element Method (BEM) head model. Current density maps were computed using a minimum norm imaging method(17).

*Computation of wSMI and PLV at source level*

We estimated the cortical sources of our "Probe epochs". We then extracted trial-to-trial source activity time series of a set of 68 cortical regions, according to a standard anatomical atlas (Desikan-Killiany(18)). Using the same method described for sensor space analysis, we computed the wSMI and the PLV between each pair of cortical regions (n=2278). Finally, to reduce the number of multiple comparisons and ease the interpretation of this analysis, we computed the median wSMI between pairs of regions of interest (ROIs). To do so, we grouped the 68 cortical regions in 5 pairs of ROIs, according to the Desikan-Killiany atlas (frontal, limbic, temporal, parietal and occipital). The wSMI/PLV between two ROIs was computed as the median across all the connections shared by the two ROIs. We hence computed a wSMI or PLV value for each pair of ROIs (n=45 connections) and each trial, at each frequency band for the wSMI (delta, theta, alpha, beta and gamma).

*Source estimation of stimulus evoked activity*

We also estimated the cortical sources of our "Target epochs". Independently for each condition (mind state), we compared the stimulus evoked activity to baseline activity (at source level) using a t-test against the baseline period (see Statistics section).

#### Statistics

Analyses are listed by order of apparition in the results section

##### *Behavior*

We compared the rate of Misses and FA between mind states at trial level using binomial generalized linear mixed models, with mind state as the explanatory factor and with two random factors (random intercepts): **subject ID and dataset (Monash dataset and Paris Brain Institute dataset**). For each behavioral measure and each mind state, response probability and confidence intervals (95%) were back-transformed from the logit scale. For

RTs, we used a (gaussian) linear mixed model with mind state as the explanatory factor and s**ubject ID and dataset as random intercepts.** Subject ID is nested within dataset as each subject belongs to only one dataset. Estimated marginal means and confidence intervals (95%) of RTs for each mind state were extracted from the model. All p-values were corrected for multiple comparisons using a *False Discovery Rate* (FDR) procedure, with an alpha level at 0.05 (5%) (for n= 24 comparisons) .

##### *Sensor space EEG markers*

For each previously described EEG marker, we compared the three mind states (ON, MW, MB) using linear mixed models at trial level, with mind state, EEG sensor (n=64) and their interaction as explanatory factors, and two random factors (random intercepts), subject identity and dataset (Monash dataset and Paris Brain Institute dataset). Marker values were standardized (z-scored) across all trials before statistical analysis to better fit the linear mixed models. ANOVA tables were computed from the models. We computed post-hoc contrasts between mind states (model estimates and statistical comparisons) at each EEG sensor. During this post-hoc analysis, to reduce the effect of possibly spurious outliers, we applied a winsorizing procedure to all contrast estimates and uncorrected p-values, were the contrast minima and maxima, as well as the p-value minima were limited to values between -5 and +5 T-values. All p-values were corrected for multiple comparisons using an FDR procedure. Main factor's p-values were corrected across the number of factors and the total number of EEG statistical models (n_factors x n_models= 3x (10 state markers + 4 source space connectivity measured + 10 ERPs time windows = 3 x 24 = 72 comparisons, alpha level = 0.05). Post-hoc p-values for each EEG marker were corrected after winsorization across the number of EEG sensors and the number of conditions (correction for n_sensors x n_conditions= 64 x 3=192 comparisons, alpha level = 0.05) .

As a control analysis, we also computed statistical models including the vigilance score, and its interaction with electrode location, as covariates, all other factors and parameters being kept identical as previously described.

##### *Source space wSMI*

For each frequency band, we performed a linear mixed model at trial level, with mind state, connection id (ROI1-ROI2) (n=45) and their interaction as explanatory factors, and the same random intercepts (subject identity and dataset). As before, we computed ANOVA tables and post-hoc contrasts between mind-states for each pair of source ROIs, correcting all p-values for multiple comparisons using an FDR procedure, as described above (correction for 72 comparisons for models' main factors, and for n_conditions x n_connections = 3 x 45 = 135 comparisons for each connectivity metric post-hoc p-values)

As a control analysis, we also computed statistical models including the vigilance score, and its interaction with connection identity, as covariates, all other factors and parameters being kept identical as previously described

##### *Sensor space ERPs*

We performed a mass-univariate (across time and space dimensions) single-trial multi-level analysis of ERPs responses. We initially intended to add both *EEG channel* and *time* (in addition to *mind state*) as factors to the same mixed linear model, but this was not feasible for computational reasons. We therefore subsampled our time dimension (-250 to 750 ms relative to stimuli onsets) by using a -25 to +25ms sliding time-window, with 75ms steps, resulting in a set of 10 time-windows of interest. We then performed a linear mixed model for each time-window, with *channel amplitude* as the dependent factor; *mind state*, *EEG sensor* and their interaction as explanatory factors; and subject ID and dataset as a random factors (random intercepts). Models' main factors' p-values were corrected for the total number of EEG statistical models x number of factors = 72 comparisons (FDR procedure, alpha level= 0.05). For each time-window, voltage estimates for each mind state, and their statistical comparisons, were computed for each EEG sensor. All obtained p-values were corrected for multiple comparisons using an FDR procedure (for n_sensors x n_conditions x n_times = 64 x 3 x 10 = 1920 comparisons, alpha level = 0.05).

##### *Temporal decoding*

For each mind-state, classification performance at each time point was tested against chance level performance using a two-sided non-parametric test (Mann-Whitney) across validation folds (distribution of scores across folds compared to a dummy beta distribution centered around 0.5, with alpha = beta= 20). These statistics were then corrected for multiple comparisons using an FDR procedure (for n_times X n_conditions= 251 x 3= 753 comparisons, alpha level = 0.05).

##### *Source estimation of stimulus evoked activity*

We performed a t-test against baseline at each time sample and each cortical source, and corrected for multiple comparisons across time, space and frequency dimensions using an FDR procedure (alpha=0.0001).

*Comparison to chance level performance of the classifier trained to distinguish different mind states*

As already explained in the previous sections, we compared our classifiers' performances to chance level using a 500-permutation procedure. We performed this procedure for each participant. At each permutation, trial labels were randomly shuffled (grouped by experimental blocks), then the entire training/testing cross-validation procedure was repeated, using the hyperparameters previously selected during the grid search procedure. We thus obtained a distribution of chance-level scores, for each participant. To calculate the individual p-value, we counted the number of permutation scores equal or higher to our true score and divided it by the number of permutations plus one. To assess group level statistical significance against chance, we performed a two-tailed Wilcoxon rank sum test, where we compared the distribution of true classification scores across participants (non permuted data), to the distribution of mean permutation scores (permuted data) across participants.

#### Software

Most analyses were performed in Python 3.9.13 , using the following packages:

- MNE (v 1.4.2) (https://mne.tools/stable/index.html): preprocessing and manipulation of EEG data
- MNE-features (v 0.2.1) (https://mne.tools/mne-features/) and nice-tools (https://github.com/nice-tools/nice): computation of EEG neural markers
- Pymer4 (v 0.8.0) (https://eshinjolly.com/pymer4/) and pingouin (v 0.5.3) (https://pingouin-stats.org/build/html/index.html) : statistical analysis
- Scikit-learn (v 1.3) (https://scikit-learn.org/stable/) : machine learning analyses

Binomial mixed models (for behavioral analyses) were performed in R using the *lme4* and *emmeans* libraries.

Source reconstruction analyses were performed in Matlab using the Brainstorm package (https://neuroimage.usc.edu/brainstorm/Introduction)

For the Monash dataset, initial preprocessing of raw EEG files using ICA was performed in Matlab using EEGlab 2021.1 and FieldTrip-20210807.

#### SUPPLEMENTARY RESULTS

#### FIGURES

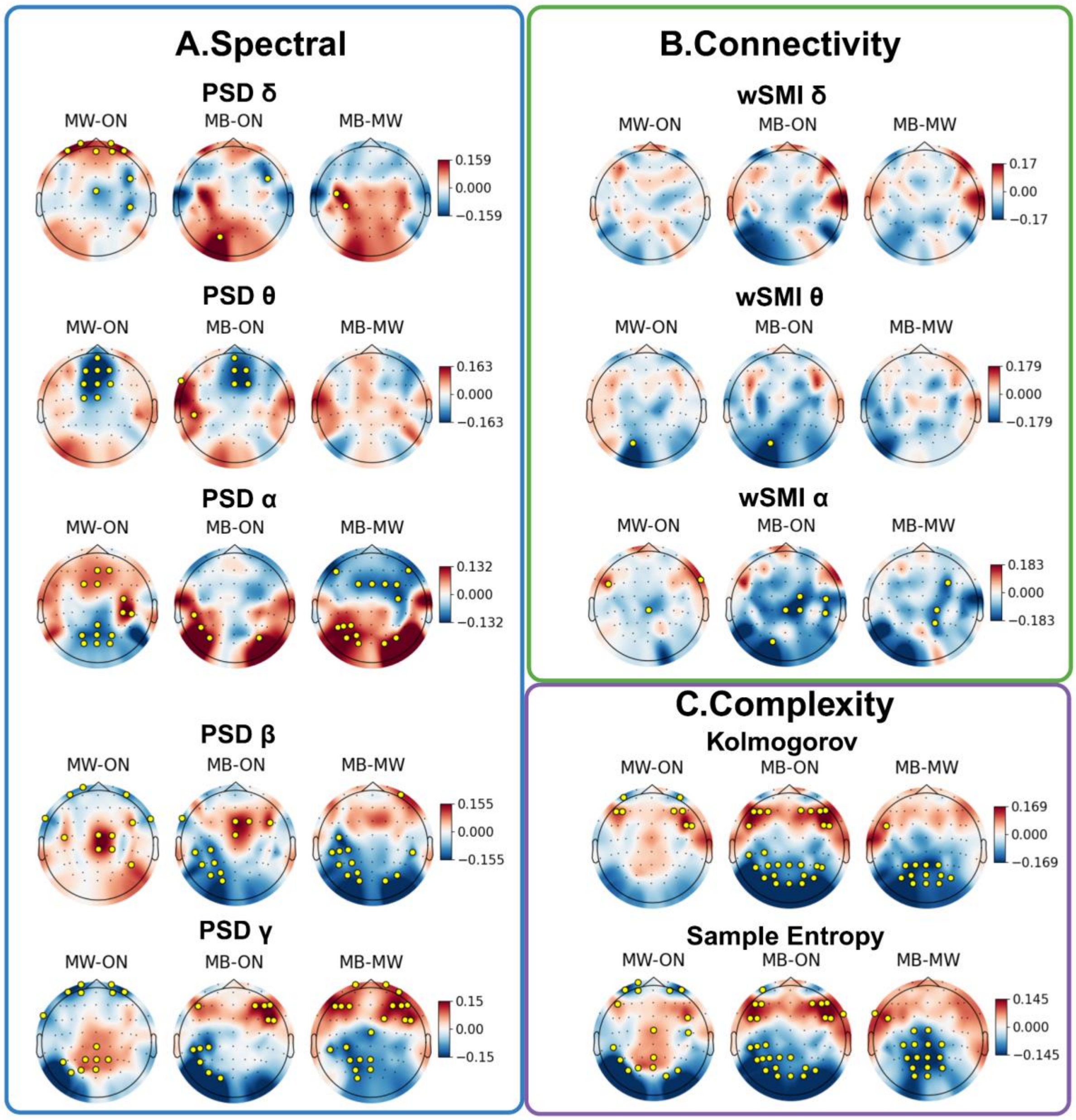


**Figure S1: Modulation of EEG markers by mind-state corrected for vigilance modifications.** Spectral (A), Connectivity (B) and Complexity (C) results. Each subplot represents the pairwise statistical contrast between two states (left: MW - ON; middle: MB - ON; right: MB - MW), at sensor level. Statistical models included the main effect of mind - state, electrode location and vigilance score (4-point scale, 1 = Extremely Sleepy, 4 = Extremely Alert), as well as the interactions mind-state/electrode location and vigilance score/electrode location. Subject ID and dataset were included as random intercepts. Model estimates for the contrast between states were computed for each electrode, and locations with statistically significant differences (FDR corrected p-value<0.05) are depicted with a golden circle. The observed results reproduced closely the results obtained with models not including vigilance score as a covariate (see main Figure 2).

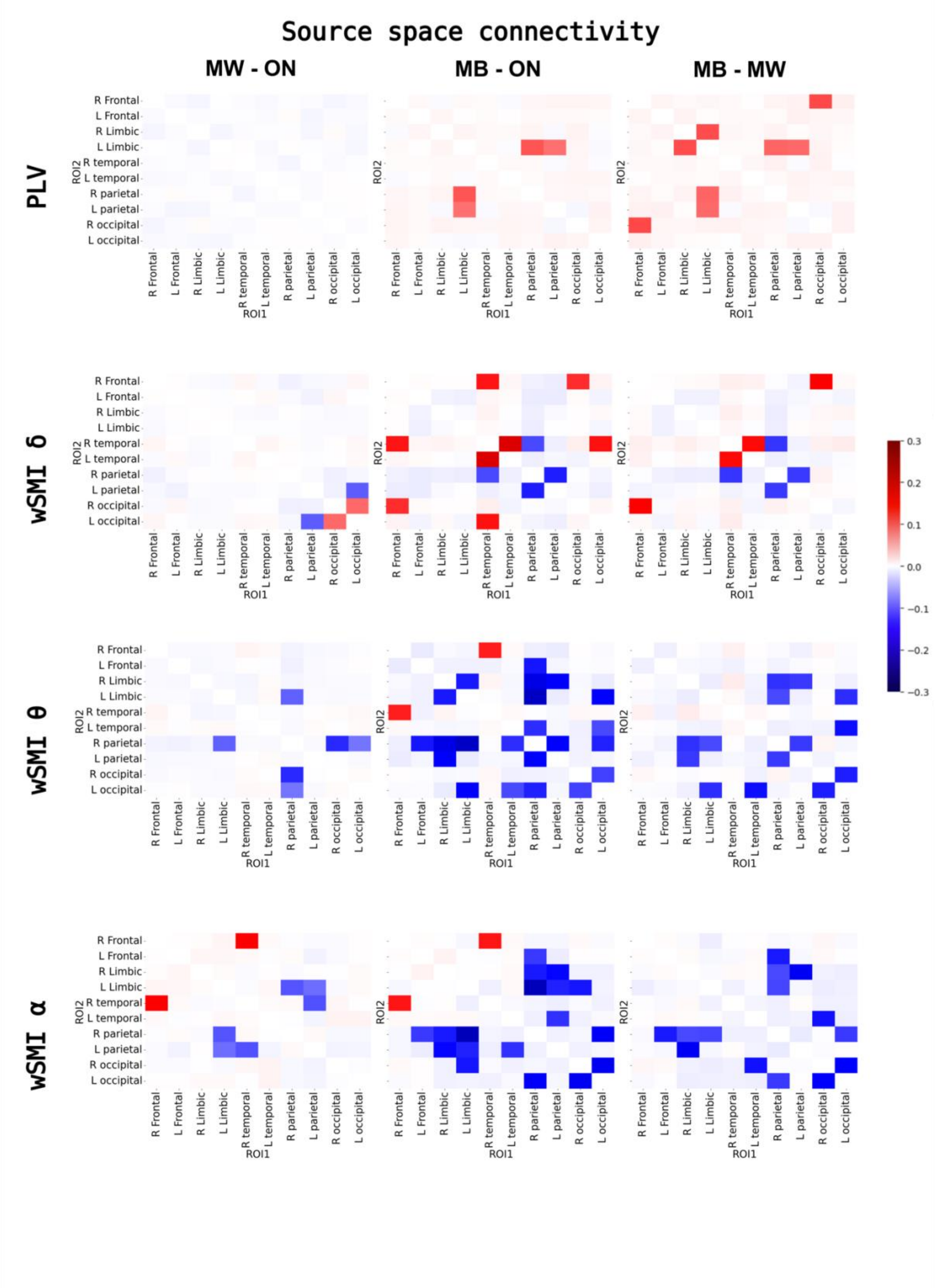


**Figure S2: Modulation of source space connectivity by mind-state corrected for vigilance modifications.** The PLV (top) and the wSMI in different frequency bands were computed at the source level. Each square-matrix represents the contrast in connectivity, computed in source space, between two mind-states, for each pair of regions of interest (ROIs). Only significantly different connections (FDR corrected p-value<0.05) are highlighted, the other ones are masked. Statistical models included the main effect of mind -state, electrode location and vigilance score (4-point scale, 1 = Extremely Sleepy, 4 = Extremely Alert), as well as the interactions mind-state/electrode location and vigilance score/electrode location. Subject ID and dataset were included as random intercepts We observed a dissociation between the PLV and wSMI, with increased PLV (phase synchrony) and decreased wSMI (information sharing) between distant cortical areas, in particular in the theta and alpha bands, during MB, as compared to ON and MW. These results are similar to those obtained without including the vigilance score in the statistical models (see Figure 3).

δ: delta band; θ: theta band; α: alpha band
wSMI: weighted symbolic mutual information
PLV: Phase Locking Value
R: right; L: left

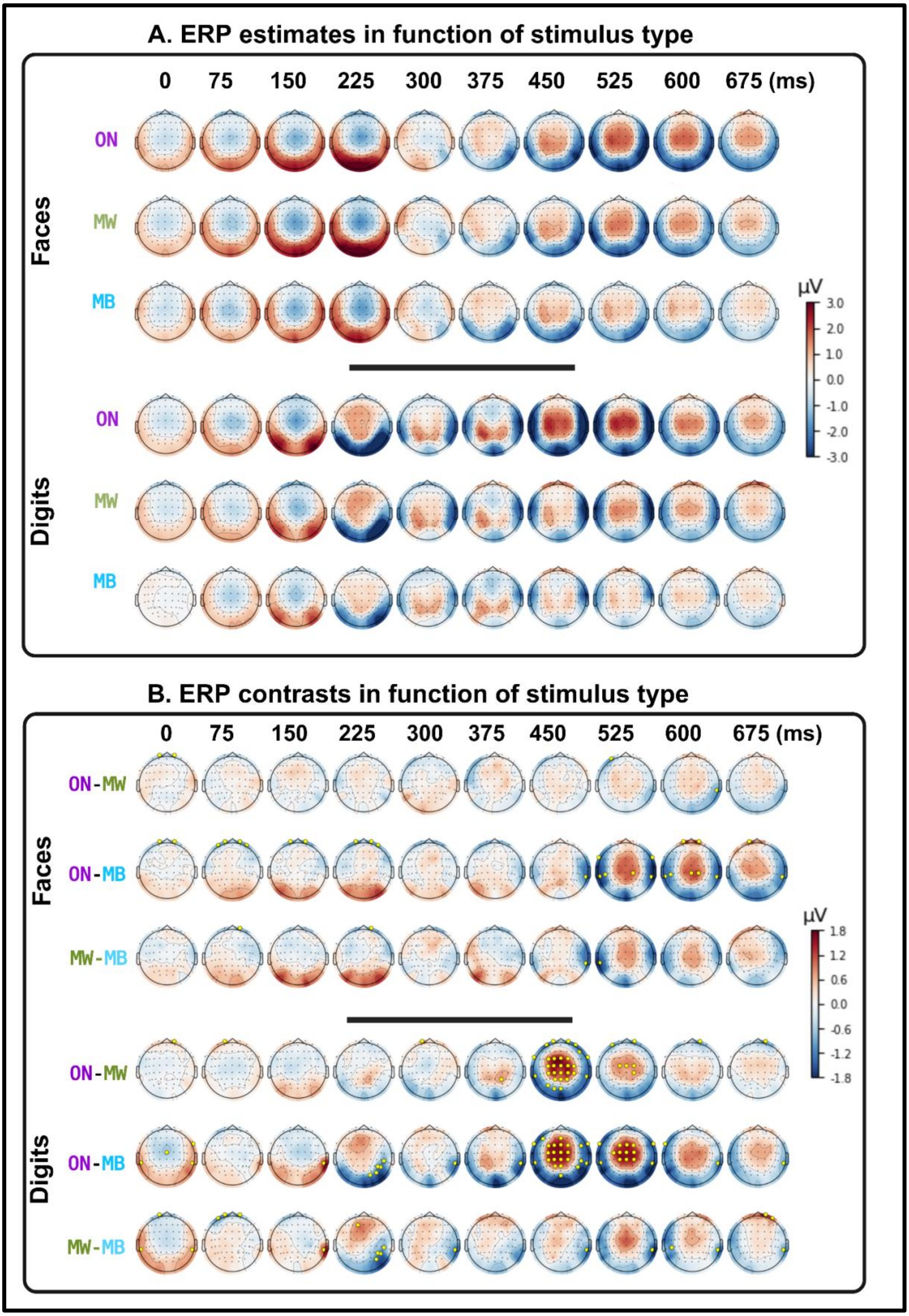
A. ERP estimates in function of stimulus type
0
75
150
225
300
375
450
525
600
675 (ms)
Faces
ON
MW
MB
Digits
ON
MW
MB
µV
3.0
2.0
1.0
0.0
-1.0
-2.0
-3.0
B. ERP contrasts in function of stimulus type
0
75
150
225
300
375
450
525
600
675 (ms)
Faces
ON-MW
ON-MB
MW-MB
Digits
ON-MW
ON-MB
MW-MB
µV
1.8
1.2
0.6
0.0
-0.6
-1.2
-1.8

**Figure S3: ERP results independently for faces and digits.** Panels A and B: Topographical scalp representation of the ERP estimates for each mind state (A) and ERP contrasts between mind-states (B), derived from the statistical model (single-trial multi-level analysis with mind-state, EEG channel and their interaction as main factors, and subject ID/dataset as random intercepts). Marked electrodes in panel B (golden circles) are the ones presenting a statistically significant difference between conditions (FDR corrected p-value<0.05). As seen in the figure, the analysis was conducted independently for faces and digits. ERPs ' profiles in function of the reported mind-state were consistent across stimuli types.

### TABLES

| Mental state | FA (%) | Misses (%) | RTs (ms) |
|---|---|---|---|
| ON | 24.5 ± 2.0 | 10.1 ± 0.9 | 471 ± 10 |
| MW | 46.1 ± 2.7 | 12.4 ± 1.1 | 465 ± 10 |
| MB | 39.9 ± 3.7 | 16.5 ± 1.6 | 480 ± 11 |
| **Behavioral measure** | **Main effect of mind state** | | **Pairwise contrasts** |
| | ***χ²*** | ***FDR corrected p-value*** | |
| FA | χ² (2)=69.8 | $8.3 * 10^{-15}$ | MW *vs* ON :<br>z-ratio=8.2,<br>p-value= $5.0 * 10^{-15}$<br><br>MB *vs* ON:<br>z-ratio=4.5,<br>p-value= $2.5 * 10^{-5}$<br><br>MB vs MW:<br>z-ratio=-1.6<br>p-value=0.12 |
| Misses | χ² (2)=64.0 | $7.6 * 10^{-14}$ | MW *vs* ON :<br>z-ratio=3.9,<br>p-value= $2.4 * 10^{-4}$<br><br>MB *vs* ON:<br>z-ratio=8.0,<br>p-value= $1.8 * 10^{-14}$<br><br>MB vs MW:<br>z-ratio=4.6<br>p-value=$1.7 * 10^{-5}$ |
| RT | F (2)=5.2 | $9.4 * 10^{-3}$ | MW *vs* ON :<br>z-ratio=-1.67,<br>p-value= 0.1<br><br>MB *vs* ON:<br>z-ratio=2.1,<br>p-value= 0.043<br><br>MB vs MW:<br>z-ratio=3.2<br>p-value=$2.6 * 10^{-3}$ |

**Table S1: Behavioral results in labeled trials.** Results of the linear mixed models with mind-state as a main factor and subject ID/dataset as random intercepts, for false alarms (FA), misses and response times (RT). The table presents the model estimates for each mind-state (top) and the statistical contrasts (bottom). All p-values were corrected for multiple comparisons using an FDR procedure.

| EEG metric | Mind -state | | Channel location | | Mind-state:Channel | |
|---|---|---|---|---|---|---|
| | F (2) | p-value | F (63) | p-value | F (126) | p -value |
| **Kolmogorov** | 19.8 | 3.78E-09 | 356.7 | 0 | 6.3 | 1.5E-97 |
| **Sample Entropy** | 46.7 | 9.49E-21 | 747.3 | 0 | 9.6 | 2.5E-175 |
| **normalized PSD delta** | 5.9 | 0.003542 | 99.6 | 0 | 3.1 | 3.5E-28 |
| **normalized PSD theta** | 8.34 | 0.00031 | 503.1 | 0 | 9.3 | 4.0E-169 |
| **normalized PSD alpha** | 62.3 | 1.73E-27 | 1335.2 | 0 | 6.5 | 4.3E-101 |
| **normalized PSD beta** | 8.3 | 0.000318 | 235.9 | 0 | 7.0 | 1.6E-113 |
| **normalized PSD gamma** | 9.4 | 0.000109 | 1180.2 | 0 | 10.0 | 1.5E-186 |
| **wSMI delta** | 37.4 | 1.03E-16 | 50.6 | 0 | 0.9 | 0.7 |
| **wSMI theta** | 57.9 | 1.37E-25 | 51.8 | 0 | 1.0 | 0.4 |
| **wSMI alpha** | 77.5 | 4.74E-34 | 69.3 | 0 | 1.9 | 1.1E-08 |

**Table S2: Neural markers results (sensor space).** ANOVA tables of the statistical models computed for each neural marker in sensor space (linear mixed models with mind state, electrode location and their interaction as main factors, and subject ID/dataset as a random intercepts). P-values were corrected for multiple comparisons using an FDR procedure.

| **Connectivity metric** | **Mind-state** | | **Connection** | | **Mind-state:Connection** | |
|---|---|---|---|---|---|---|
| | F (2) | p-value | F (44) | p-value | F (88) | p-value |
| **wSMI alpha** | 32.9 | 8.52E-15 | 352.5 | 0 | 1.7 | 0.0001 |
| **wSMI theta** | 42.9 | 3.95E-19 | 213.3 | 0 | 1.1 | 0.2 |
| **wSMI delta** | 24.8 | 2.9E-11 | 203.8 | 0 | 1.2 | 0.1 |

**Table S3: Source-level connectivity.** ANOVA tables of the statistical models computed for each connectivity measure in source space (linear mixed models with mind state, connection ID and their interaction as main factors, and subject ID/dataset as random intercepts). P-values were corrected for multiple comparisons using an FDR procedure.

| **Time-window (-/+ 25ms)** | **Mind-state** | | **Channel location** | | **Mind-state:Channel location** | |
|---|---|---|---|---|---|---|
| | F (2) | p-value | F (63) | p-value | F (126) | p-value |
| **0 ms** | 0.3 | 0.8 | 48.9 | 0 | 1.9 | 5.3E-10 |
| **75 ms** | 0.5 | 0.7 | 83.2 | 0 | 1.9 | 8.0E-10 |
| **150 ms** | 0.3 | 0.8 | 138.9 | 0 | 1.5 | 0.0003 |
| **225 ms** | 0.2 | 0.9 | 32.0 | 0 | 1.8 | 3.1E-07 |
| **300 ms** | 0.2 | 0.9 | 34.5 | 0 | 1.1 | 0.4 |
| **375 ms** | 0.07 | 0.9 | 38.9 | 0 | 1.8 | 3.4E-08 |
| **450 ms** | 0.2 | 0.9 | 91.9 | 0 | 6.6 | 8.6E-103 |
| **525 ms** | 0.5 | 0.7 | 119.9 | 0 | 7.7 | 2.1E-130 |
| **600 ms** | 0.6 | 0.6 | 83.8 | 0 | 4.4 | 1.6E-55 |
| **675 ms** | 0.3 | 0.8 | 52.4 | 0 | 2.7 | 1.5E-21 |

**Table S4 : Event related potentials (ERPs).** ANOVA tables of the statistical models computed at each time window (linear mixed models with mind state, electrode location and their interaction as main factors, and subject ID/dataset as random intercepts). P-values were corrected for multiple comparisons using an FDR procedure.

| | ON | MW | MB |
|---|---|---|---|
| **Precision** | 0.598 | 0.498 | 0.278 |
| **Recall** | 0.631 | 0.516 | 0.309 |
| **F1-score** | 0.605 | 0.498 | 0.289 |

**Table S5:** Weighted-average performance of individual's classifiers (PCA/Random Forest classifier) for each mind-state. The metrics (precision, recall, f1-score) are computed as the weighted mean across subjects, where each subject's contribution is weighted by the number of trials (support) for the respective class.

| | MB | | | MW | | | ON | | | Balanced accuracy/permutation test | | |
|---|---|---|---|---|---|---|---|---|---|---|---|---|
| **Subject** | **Pr** | **Re** | **F1** | **Pr** | **Re** | **F1** | **Pr** | **Re** | **F1** | **Observed** | **Perm mean** | **p-value** |
| Monash 001 | NA | NA | NA | 0.562 | 0.551 | 0.557 | 0.745 | 0.753 | 0.749 | 0.6805 | 0.5035 | 0.0040 |
| Monash 002 | NA | NA | NA | 0.316 | 0.200 | 0.245 | 0.913 | 0.951 | 0.931 | 0.7321 | 0.6503 | 0.0200 |
| **Monash 003** | **0.403** | **0.475** | **0.436** | **0.200** | **0.162** | **0.179** | **0.299** | **0.277** | **0.287** | **0.3524** | **0.3301** | **0.6387** |
| **Monash 004** | **0.392** | **0.305** | **0.343** | **0.237** | **0.136** | **0.173** | **0.475** | **0.656** | **0.551** | **0.4325** | **0.3340** | **0.0080** |
| **Monash 005** | **0.075** | **0.167** | **0.103** | **0.218** | **0.449** | **0.294** | **0.700** | **0.295** | **0.415** | **0.4505** | **0.3451** | **0.1018** |
| Monash 006 | NA | NA | NA | 0.794 | 0.992 | 0.882 | 0.000 | 0.000 | 0.000 | 0.5943 | 0.5961 | 0.8563 |
| **Monash 007** | **0.133** | **0.053** | **0.075** | **0.511** | **0.627** | **0.563** | **0.291** | **0.255** | **0.272** | **0.4134** | **0.3917** | **0.8822** |
| **Monash 008** | **0.115** | **0.111** | **0.113** | **0.486** | **0.493** | **0.490** | **0.427** | **0.424** | **0.426** | **0.4336** | **0.3583** | **0.1058** |
| **Monash 009** | **0.432** | **0.488** | **0.458** | **0.476** | **0.291** | **0.361** | **0.155** | **0.391** | **0.222** | **0.4272** | **0.3503** | **0.0259** |
| **Monash 010** | **0.244** | **0.270** | **0.256** | **0.450** | **0.386** | **0.415** | **0.426** | **0.433** | **0.430** | **0.4103** | **0.3322** | **0.0758** |
| Monash 011 | NA | NA | NA | 0.000 | 0.000 | 0.000 | 0.845 | 0.992 | 0.912 | 0.6955 | 0.6986 | 0.9082 |
| **Monash 012** | **0.264** | **0.383** | **0.313** | **0.569** | **0.412** | **0.478** | **0.298** | **0.364** | **0.327** | **0.4037** | **0.3387** | **0.3353** |

| | | | | | | | | | | | | |
|---|---|---|---|---|---|---|---|---|---|---|---|---|
| Monash 013 | NA | NA | NA | NA | NA | NA | 1.000 | 1.000 | 1.000 | 1.000 | 1.000 | 1.0 |
| **Monash 014** | **0.000** | **0.000** | **0.000** | **0.174** | **0.043** | **0.069** | **0.634** | **0.899** | **0.744** | **0.5769** | **0.5442** | **0.2255** |
| **Monash 015** | **0.000** | **0.000** | **0.000** | **0.531** | **0.673** | **0.594** | **0.423** | **0.275** | **0.333** | **0.4281** | **0.3855** | **0.6327** |
| **Monash 016** | **0.232** | **0.333** | **0.274** | **0.429** | **0.368** | **0.396** | **0.282** | **0.296** | **0.289** | **0.3891** | **0.3428** | **0.4471** |
| **Monash 017** | **0.059** | **0.046** | **0.052** | **0.298** | **0.330** | **0.313** | **0.533** | **0.550** | **0.541** | **0.3408** | **0.3408** | **0.6248** |
| **Monash 018** | **0.000** | **0.000** | **0.000** | **0.678** | **0.775** | **0.723** | **0.100** | **0.070** | **0.083** | **0.4963** | **0.4604** | **0.6906** |
| **Monash 019** | **0.503** | **0.593** | **0.545** | **0.239** | **0.145** | **0.180** | **0.323** | **0.352** | **0.337** | **0.4368** | **0.3373** | **0.2076** |
| **Monash 020** | **0.000** | **0.000** | **0.000** | **0.135** | **0.086** | **0.105** | **0.757** | **0.864** | **0.807** | **0.4522** | **0.4338** | **0.4691** |
| **Monash 021** | **0.000** | **0.000** | **0.000** | **0.086** | **0.049** | **0.062** | **0.710** | **0.850** | **0.774** | **0.5463** | **0.5257** | **0.8204** |
| **Monash 022** | **0.000** | **0.000** | **0.000** | **0.547** | **0.611** | **0.577** | **0.552** | **0.540** | **0.546** | **0.5877** | **0.4632** | **0.2056** |
| **Monash 023** | **0.000** | **0.000** | **0.000** | **0.423** | **0.294** | **0.347** | **0.642** | **0.782** | **0.705** | **0.5031** | **0.4343** | **0.0878** |
| **Monash 024** | **0.000** | **0.000** | **0.000** | **0.359** | **0.137** | **0.199** | **0.540** | **0.859** | **0.663** | **0.3831** | **0.3305** | **0.0399** |
| Monash 025 | NA | NA | NA | 0.783 | 0.864 | 0.822 | 0.348 | 0.232 | 0.278 | 0.6912 | 0.5992 | 0.0259 |
| **Monash 026** | **0.000** | **0.000** | **0.000** | **0.778** | **0.871** | **0.822** | **0.420** | **0.300** | **0.350** | **0.5712** | **0.5423** | **0.1677** |
| **Paris 001** | **0.211** | **0.154** | **0.178** | **0.521** | **0.656** | **0.580** | **0.393** | **0.276** | **0.324** | **0.4264** | **0.3355** | **0.0299** |
| **Paris 002** | **0.480** | **0.571** | **0.522** | **0.316** | **0.276** | **0.294** | **0.385** | **0.321** | **0.350** | **0.4377** | **0.3528** | **0.3154** |

| | | | | | | | | | | | |
|---|---|---|---|---|---|---|---|---|---|---|---|
| Paris 003 | 0.389 | 0.323 | 0.353 | 0.637 | 0.745 | 0.687 | 0.148 | 0.085 | 0.108 | 0.4181 | 0.3615 | 0.1617 |
| Paris 004 | 0.130 | 0.062 | 0.085 | 0.366 | 0.293 | 0.325 | 0.400 | 0.540 | 0.460 | 0.4414 | 0.3783 | 0.4511 |
| Paris 005 | 0.358 | 0.375 | 0.366 | 0.463 | 0.452 | 0.458 | 0.398 | 0.398 | 0.398 | 0.4325 | 0.3374 | 0.0200 |
| Paris 006 | 0.000 | 0.000 | 0.000 | 0.382 | 0.366 | 0.374 | 0.647 | 0.776 | 0.706 | 0.4394 | 0.3603 | 0.0479 |
| Paris 007 | 0.000 | 0.000 | 0.000 | 0.520 | 0.552 | 0.536 | 0.700 | 0.684 | 0.692 | 0.5831 | 0.4713 | 0.0160 |
| Paris 008 | 0.161 | 0.310 | 0.212 | 0.388 | 0.392 | 0.390 | 0.655 | 0.547 | 0.596 | 0.5069 | 0.3629 | 0.0020 |
| Paris 009 | 0.309 | 0.630 | 0.415 | 0.312 | 0.164 | 0.215 | 0.721 | 0.613 | 0.663 | 0.4295 | 0.3346 | 0.0060 |
| Paris 010 | 0.200 | 0.222 | 0.211 | 0.443 | 0.360 | 0.397 | 0.678 | 0.729 | 0.703 | 0.4759 | 0.3574 | 0.0020 |
| Paris 011 | 0.000 | 0.000 | 0.000 | 0.125 | 0.017 | 0.030 | 0.629 | 0.914 | 0.745 | 0.5198 | 0.5191 | 0.7066 |
| Paris 012 | 0.333 | 0.419 | 0.371 | 0.511 | 0.463 | 0.486 | 0.146 | 0.125 | 0.135 | 0.4235 | 0.3422 | 0.0958 |
| Paris 013 | 0.461 | 0.519 | 0.488 | 0.000 | 0.000 | 0.000 | 0.611 | 0.725 | 0.663 | 0.4728 | 0.3503 | 0.0060 |
| Paris 014 | 0.125 | 0.256 | 0.168 | 0.106 | 0.292 | 0.156 | 0.853 | 0.540 | 0.661 | 0.4463 | 0.3560 | 0.2774 |
| Paris 015 | 0.000 | 0.000 | 0.000 | 0.351 | 0.283 | 0.313 | 0.837 | 0.884 | 0.860 | 0.5066 | 0.4472 | 0.0080 |
| Paris 016 | 0.000 | 0.000 | 0.000 | 0.444 | 0.330 | 0.379 | 0.709 | 0.814 | 0.758 | 0.5167 | 0.4674 | 0.1198 |
| Paris 017 | 0.354 | 0.347 | 0.350 | 0.118 | 0.095 | 0.105 | 0.589 | 0.628 | 0.608 | 0.4727 | 0.3302 | 0.0100 |
| Paris 018 | 0.000 | 0.000 | 0.000 | 0.000 | 0.000 | 0.000 | 0.887 | 1.000 | 0.940 | 0.6767 | 0.6642 | 0.0359 |

| | | | | | | | | | | | |
|---|---|---|---|---|---|---|---|---|---|---|---|
| **Paris 019** | **0.000** | **0.000** | **0.000** | **0.000** | **0.000** | **0.000** | **0.746** | **0.946** | **0.834** | **0.4836** | **0.4969** | **1.0** |
| **Paris 020** | **0.000** | **0.000** | **0.000** | **0.250** | **0.328** | **0.284** | **0.783** | **0.731** | **0.756** | **0.5917** | **0.5353** | **0.1796** |
| Paris 021 | NA | NA | NA | 0.269 | 0.206 | 0.233 | 0.773 | 0.829 | 0.800 | 0.5267 | 0.4979 | 0.4311 |
| **Paris 022** | **0.036** | **0.071** | **0.048** | **0.497** | **0.657** | **0.566** | **0.375** | **0.206** | **0.266** | **0.4624** | **0.3788** | **0.1876** |
| **Paris 023** | **0.000** | **0.000** | **0.000** | **0.000** | **0.000** | **0.000** | **0.600** | **0.907** | **0.722** | **0.4985** | **0.5135** | **0.7485** |
| **Paris 024** | **0.550** | **0.484** | **0.515** | **0.000** | **0.000** | **0.000** | **0.611** | **0.739** | **0.669** | **0.5353** | **0.4324** | **0.0160** |
| **Paris 025** | **0.447** | **0.273** | **0.339** | **0.350** | **0.500** | **0.412** | **0.377** | **0.460** | **0.414** | **0.4773** | **0.3320** | **0.0020** |
| **Paris 026** | **0.000** | **0.000** | **0.000** | **0.000** | **0.000** | **0.000** | **0.817** | **0.992** | **0.896** | **0.4333** | **0.4324** | **0.6567** |
| **Paris 027** | **0.000** | **0.000** | **0.000** | **0.579** | **0.596** | **0.587** | **0.627** | **0.631** | **0.629** | **0.5789** | **0.4578** | **0.0020** |
| **Paris 028** | **0.382** | **0.317** | **0.347** | **0.465** | **0.444** | **0.454** | **0.500** | **0.555** | **0.526** | **0.4328** | **0.3821** | **0.3253** |
| **Paris 029** | **0.000** | **0.000** | **0.000** | **0.590** | **0.727** | **0.651** | **0.432** | **0.312** | **0.363** | **0.4850** | **0.4429** | **0.0978** |
| **Paris 030** | **0.218** | **0.250** | **0.233** | **0.472** | **0.402** | **0.434** | **0.431** | **0.470** | **0.450** | **0.4202** | **0.2885** | **0.0020** |
| **Paris 031** | **0.091** | **0.042** | **0.057** | **0.270** | **0.233** | **0.250** | **0.687** | **0.768** | **0.725** | **0.4677** | **0.3857** | **0.0240** |
| Paris 032 | NA | NA | NA | NA | NA | NA | 1.000 | 1.000 | 1.000 | 1.000 | 1.000 | 1.0 |
| **Paris 033** | **0.273** | **0.284** | **0.278** | **0.233** | **0.206** | **0.219** | **0.538** | **0.556** | **0.547** | **0.3660** | **0.3457** | **0.2176** |
| **Paris 034** | **0.000** | **0.000** | **0.000** | **0.462** | **0.362** | **0.406** | **0.621** | **0.728** | **0.670** | **0.5940** | **0.5215** | **0.1178** |

| | | | | | | | | | | | |
|---|---|---|---|---|---|---|---|---|---|---|---|
| **Paris 035** | **0.000** | **0.000** | **0.000** | **0.500** | **0.667** | **0.571** | **0.415** | **0.300** | **0.348** | **0.4877** | **0.4284** | **0.2475** |
| **Paris 036** | **0.222** | **0.229** | **0.225** | **0.315** | **0.299** | **0.307** | **0.560** | **0.574** | **0.567** | **0.3897** | **0.3309** | **0.0060** |

**Table S6: Detailed results of subject level classifiers.** For each subject and each class, precision (Pr), recall (Re) and F-1 score are presented. For each subject, the observed balanced accuracy score, the mean balanced accuracy of the permutation test and the p-value of the permutation test are presented. Highlighted subjects (in bold) present with the 3 classes (3 mind states) and were included in the group analysis (Figure 6, n= 54/62)

| Predicted Mental state | FA (%) | Misses (%) | RTs (ms) |
|---|---|---|---|
| ON | 32.1 ± 2.9 | 1.3 ± 0.4 | 468 ± 10 |
| MW | 39.2 ± 3.5 | 1.8 ± 0.6 | 475 ± 10 |
| MB | 37.6 ± 4.2 | 2.8 ± 0.9 | 483 ± 11 |
| **Behavioral measure** | **Main effect of mind state** | | **Pairwise contrasts** |
| | ***χ²*** | ***FDR corrected p-value*** | |
| FA | $\chi^2$ (2)=6.8 | 0.04 | MW *vs* ON :<br>z-ratio=2.5, p-value= 0.02<br>MB *vs* ON:<br>z-ratio=1.5, p-value= 0.1<br>MB vs MW:<br>z-ratio=-0.4, p-value=0.7 |
| Misses | $\chi^2$ (2)=31.5 | $5.7 * 10^{-7}$ | MW *vs* ON :<br>z-ratio=2.4, p-value= 0.02<br>MB *vs* ON:<br>z-ratio=5.6, p-value=$9.6 * 10^{-8}$<br>MB vs MW:<br>z-ratio=3.3, p-value=0.002 |
| RT | F (2)=7.4 | $1.4 * 10^{-3}$ | MW *vs* ON :<br>z-ratio=2.3, p-value= 0.03<br>MB *vs* ON:<br>z-ratio=3.6, p-value= $7.5 * 10^{-4}$<br>MB vs MW:<br>z-ratio=1.8 p-value=0.08 |

**Table S7: Behavioral results in non-labeled trials in function of the predicted mind-state.** Results of the linear mixed models with predicted mind-state as a main factor and subject ID/dataset as random intercepts, for false alarms (FA), misses and response times (RT). The table presents the model estimates for each mind -state (top) and the statistical contrasts (bottom). P-values were corrected for multiple comparisons using an FDR procedure.

#### SUPPLEMENTARY REFERENCES


1. O. Langner, *et al.*, Presentation and validation of the Radboud Faces Database. *Cognition & Emotion* **24**, 1377–1388 (2010).

2. C. Van den Driessche, *et al.*, Attentional Lapses in Attention-Deficit/Hyperactivity Disorder: Blank Rather Than Wandering Thoughts. *Psychol Sci* **28**, 1375–1386 (2017).

3. J.-R. King, *et al.*, Information Sharing in the Brain Indexes Consciousness in Noncommunicative Patients. *Current Biology* **23**, 1914–1919 (2013).

4. J. D. Sitt, *et al.*, Large scale screening of neural signatures of consciousness in patients in a vegetative or minimally conscious state. *Brain* **137**, 2258–2270 (2014).

5. D. A. Engemann, *et al.*, Robust EEG-based cross-site and cross-protocol classification of states of consciousness. *Brain* **141**, 3179–3192 (2018).

6. P. Bourdillon, *et al.*, Brain-scale cortico-cortical functional connectivity in the delta-theta band is a robust signature of conscious states: an intracranial and scalp EEG study. *Scientific Reports* **10**, 1–13 (2020).

7. B. Türker, *et al.*, Behavioral and brain responses to verbal stimuli reveal transient periods of cognitive integration of external world in all sleep stages. [Preprint] (2022). Available at: https://www.biorxiv.org/content/10.1101/2022.05.04.490484v1 [Accessed 4 August 2023].

8. L. S. Imperatori, *et al.*, EEG functional connectivity metrics wPLI and wSMI account for distinct types of brain functional interactions. *Sci Rep* **9**, 8894 (2019).

9. R. Martínez Vivot, C. Pallavicini, F. Zamberlan, D. Vigo, E. Tagliazucchi, Meditation Increases the Entropy of Brain Oscillatory Activity. *Neuroscience* **431**, 40–51 (2020).

10. J. Kayser, C. E. Tenke, Principal components analysis of Laplacian waveforms as a generic method for identifying ERP generator patterns: II. Adequacy of low-density estimates. *Clinical Neurophysiology* **117**, 369–380 (2006).

11. J. S. Richman, J. R. Moorman, Physiological time-series analysis using approximate entropy and sample entropy. *American Journal of Physiology-Heart and Circulatory Physiology* **278**, H2039–H2049 (2000).

12. H. I. Volpert-Esmond, E. C. Merkle, M. P. Levsen, T. A. Ito, B. D. Bartholow, Using Trial-Level Data and Multilevel Modeling to Investigate Within-Task Change in Event-Related Potentials. *Psychophysiology* **55**, e13044 (2018).

13. J.-R. King, S. Dehaene, Characterizing the dynamics of mental representations: the temporal generalization method. *Trends in Cognitive Sciences* **18**, 203–210 (2014).

14. J.-R. King, *et al.*, Encoding and Decoding Neuronal Dynamics: Methodological Framework to Uncover the Algorithms of Cognition. (2018).

15. J.-R. King, A. Gramfort, A. Schurger, L. Naccache, S. Dehaene, Two Distinct Dynamic Modes Subtend the Detection of Unexpected Sounds. *PLoS ONE* **9**, e85791 (2014).

16. J. Weston, B. Schölkopf, G. Bakir, Learning to Find Pre-Images in *Advances in Neural Information Processing Systems*, (MIT Press, 2003).

17. S. Baillet, J. C. Mosher, R. M. Leahy, Electromagnetic brain mapping. *IEEE Signal Processing Magazine* **18**, 14–30 (2001).

18. R. S. Desikan, *et al.*, An automated labeling system for subdividing the human cerebral cortex on MRI scans into gyral based regions of interest. *NeuroImage* **31**, 968–980 (2006).
